# CoRe: A robustly benchmarked R package for identifying core-fitness genes in genome-wide pooled CRISPR-Cas9 screens

**DOI:** 10.1101/2021.05.25.445610

**Authors:** Alessandro Vinceti, Emre Karakoc, Clare Pacini, Umberto Perron, Riccardo Roberto De Lucia, Mathew J. Garnett, Francesco Iorio

## Abstract

CRISPR-Cas9 genome-wide screens are being increasingly performed, allowing systematic explorations of cancer dependencies at unprecedented accuracy and scale. Identifying from these screens the genes that are essential for cell survival invariantly across tissues, conditions, and genomic-contexts (core-fitness genes), is of paramount importance to assess the safety profile of candidate therapeutic targets and for elucidating mechanisms involved in tissue-specific genetic diseases. We present CoRe: An R package implementing novel methods for identifying core-fitness genes from joint analyses of multiple CRISPR-Cas9 screens. We demonstrate that CoRe outperforms state-of-the-art tools, yielding more reliable sets of core-fitness genes than existing and widely used reference sets.

## Background

The ability to perturb individual genes at scale in human cells holds the key to elucidating their function and it is a gateway to the identification of new therapeutic targets across human diseases, including cancer. In this context the CRISPR-Cas9 genome editing system is the state-of-the-art tool [1–3].

Several genome-scale CRISPR-Cas9 single guide RNA (sgRNA) libraries have been designed and are available to date for genetic perturbation screens in human cells, showing significantly improved precision and scale with respect to previous technologies [4–8]. Some of these libraries have been employed in large-scale *in-vitro* screens assessing each gene’s potential in reducing cellular viability/fitness upon inactivation, across hundreds of immortalised human cancer cell lines [7, 9–12]. This has led to comprehensive identifications of cellular fitness genes, providing a detailed view of genetic dependencies and vulnerabilities existing in cancer cells.

Several sources of bias must be considered when analysing dependency profiles derived from CRISPR-Cas9 screens. These include different guide efficiency and off-target effects [13, 14], genomic features like copy number amplifications [7, 15–17], variable phenotypic penetrance [18], and different experimental settings such as, for example, screening time length and cells’ growth medium [19, 20]. Taken together, these factors contribute to making the analysis of CRISPR-Cas9 screens not trivial, and several tools have been proposed for this task [21–27].

When analysing data from CRISPR-Cas9 screens in functional and translational studies another major computational problem is to classify and distinguish genetic dependencies involved in normal essential biological processes from disease- and genomic-context-specific vulnerabilities.

Identifying context-specific essential genes, and distinguishing them from constitutively essential genes shared across all tissues and cells, i.e. core-fitness genes (CFGs), is also crucial for elucidating the mechanisms involved in tissue-specific diseases. Moving forward and focusing on very well-defined genomic contexts in tumours allows identifying cancer synthetic lethalities that could be exploited therapeutically [28].

Gene dependency profiles, generated via pooled CRISPR-Cas9 screening across large panels of human cancer cell lines, are becoming increasingly available [29, 30]. However, identifying and discriminating CFGs and context-specific essential genes from this type of functional genetics screens remains not trivial.

The Daisy Model (DM) has been recently described for identifying CFGs by jointly analysing data from genetic screens of multiple cancer cell lines. In this approach, sets of fitness genes for each screened cancer cell line are conceptually represented by the petals of a daisy [10]. These have different extents of overlap, but they generally tend to share a common set of CFGs (the core of the daisy). Based on this idea, genes that are essential in most of the screened cell lines are predicted to be CFGs. This approach has been shown to be able to identify CFGs that are enriched for fundamental cellular processes such as transcription, translation, and replication [10]. Nevertheless, in [10] the minimal number of cell lines (3 out of 5 screened), in which a gene should be significantly essential in order to be predicted as CFG, is arbitrarily defined with no indications on how to determine this threshold on a numerically grounded basis when applying the DM to larger collections of screens.

In [12] we have introduced the Adaptive Daisy Model (ADaM): a generalisation of the DM that is able to determine the minimal number of cell lines that should be vulnerable to knocking-out the putative CFGs, i.e. dependent on them, in a semi-supervised manner. We have also recently proposed an alternative unsupervised approach within the Broad and Sanger Institutes’ Cancer Dependency Map collaboration [31, 32], where data from screening hundreds of cell lines are analysed in a pooled fashion, independently of their tissue of origin. This method builds on the intuition that if a gene is universally essential then it should rank among the top essential genes in most screened models, including those that are the least dependent on it, or generally show a moderate to weak loss-of-fitness phenotype upon CRISPR-Cas9 targeting.

Finally, a logistic regression based method for classifying genes into CFGs or context-specific essentials has been recently introduced by Sharma and colleagues [33] as part of the CEN-tools suite, using reference sets of essential and non-essential genes for the training phase [34].

Although the number of CRISPR-Cas9 and genome-scale RNAi experiments is increasing rapidly, no robustly benchmarked method to identify sets of CFGs has been devised yet in a unique and easy-to-use software package.

We present CoRe: an R package implementing recently proposed as well as novel versions of algorithms for the identification of CFGs from a joint analysis of multiple genome-wide pooled CRISPR-Cas9 knock-out screens. Furthermore, we present results from a comparison of CoRe’s output (when applied to the largest integrative cancer dependency dataset generated to date [19]) against widely used [10, 34], or more recent [33] sets of CFGs obtained via an alternative approach (which we have also tested on the same recent cancer dependency dataset). We report an increased coverage of prior known human essential genes, new potential core-fitness genes, and lower false positive rates for CoRe’s methods with respect to other state-of-the-art core-fitness sets and available methods. Finally we show that CoRe’s methods are computationally more efficient than others, and that the CFGs obtained with CoRe could be used in the future as a template classifier of a single screen’s specific essential genes, via supervised classification methods, such as the widely used BAGEL [26].

## Results

### Overview of the CoRe package and implemented methods

We have developed and extensively benchmarked CoRe: An R package able to identify core-fitness genes (CFGs) from the joint analysis of multiple genome-wide CRISPR knock-out screens.

CoRe implements two methods at two different levels of stringency yielding, respectively, (i) CFGs and (ii) common-essential genes (CEGs). Both sets include genes that are essential for cell survival invariantly across tissues and genomic backgrounds and are involved in housekeeping cellular processes, thus are conceptually the same. However, CFGs are identified in CoRe more stringently and in a supervised manner, whereas CEGs are outputted by a less stringent and unsupervised method. These two-level of stringency make CoRe suitable for a variety of use-case scenarios. These range from the robust identification of new human core essential genes (where minimising false positive is essential, thus CFGs should be preferred to CEGs), to filtering out potential cytotoxic candidates when focusing on context-specific essential genes while identifying and prioritising new therapeutic targets (where is more important to minimise the false negatives, thus CEGs should be preferred to CFGs).

The first and more stringent method implemented in CoRe is the Adaptive Daisy Model (ADaM) [12]: an adaptive version of the Daisy Model (DM) [10] that operates in a cascade of two steps, and it is usable on data coming from large-scale CRISPR-cas9 knock-out screens performed in heterogeneous in vitro models, for example immortalized human cancer cell lines from multiple tissue lineages (**Fig. 1A-D**).

**Figure 1-.**
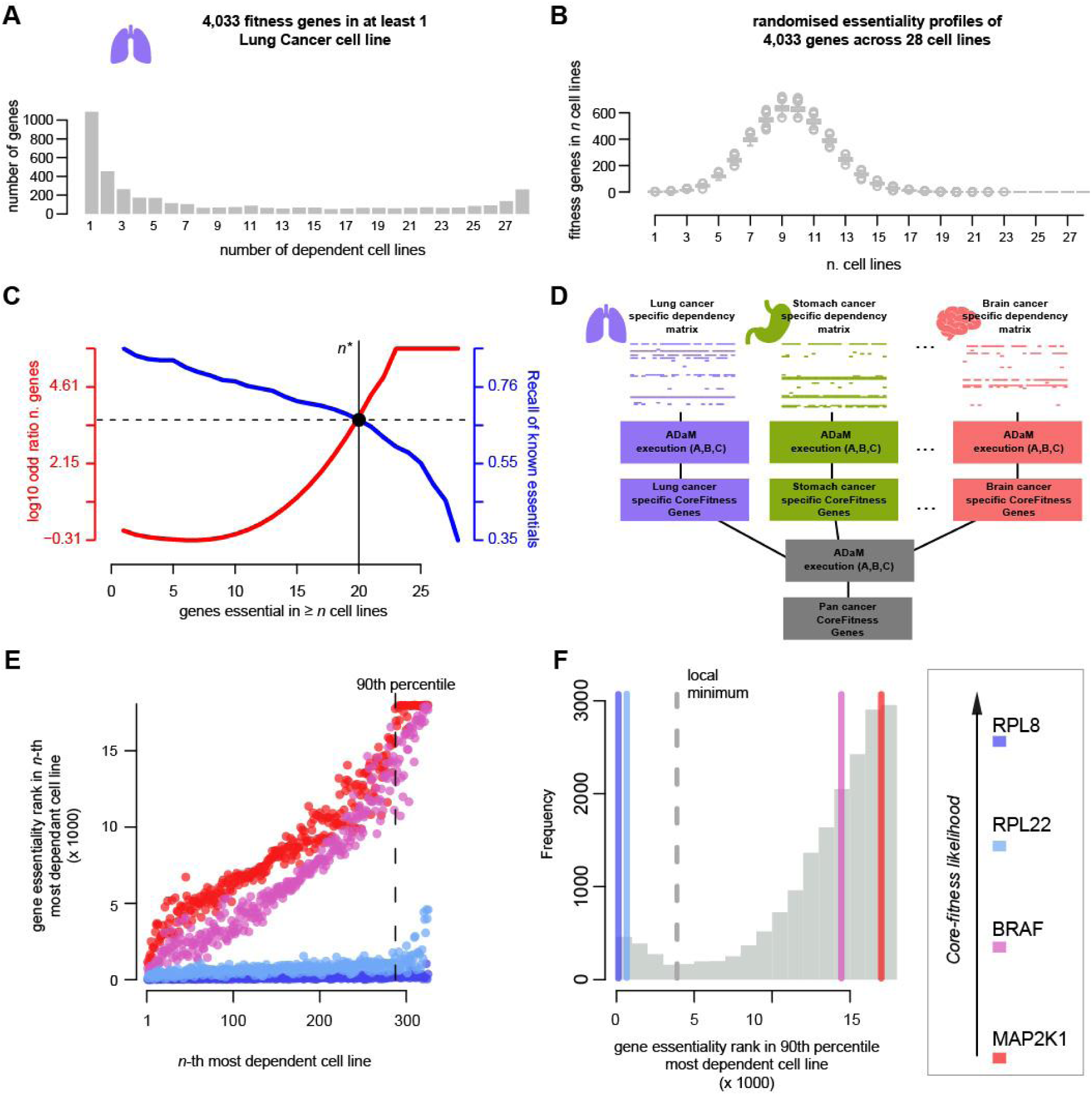
Overview of the methods implemented in CoRe. **A**. Number of fitness genes in fixed numbers of cell lines (CLs) from a lung specific binary cancer dependency matrix (BM). **B**. As for A but considering 1,000 randomisations of the lung BM. **C**. ADaM execution on the lung BM: The aim is to identify the minimal number *n*\* of CLs in which a gene should be essential to be considered a lung specific core-fitness essential gene (CFG). All possible *n* values (on the x-axis) are tested. For each *n* the genes essential in ≥ *n* CLs are determined. The Recall of a reference set of CFGs (blue curve, and right y-axis) is computed for this set of genes. At the same time the deviance of expectation of the size of this set of genes is also computed (log_10_ ratio with respect to average value in 10,000 permutations of the lung BM (red curve, and left y-axis). The *n*\* value (solid vertical line) is that providing the best trade-off (dashed horizontal line) between the blue and the red curves. **D**. Schematic of the two-step model of ADaM identifying pan-cancer CFGs. The first determines sets of tissue/cancer-type specific CFGs. The second step computes pan-cancer CFGs as those predicted as tissues/cancer-type specific core-fitness genes for at least *t*\* tissues/cancer-types. This is determined as for the *n*\* in C. E. Basic assumption of the FiPer method: common-essential genes (CEGs) are always among the top essential genes. 4 example genes are shown. Each point indicates a CL. The coordinate on the x-axis indicates the rank position of the CL when sorting all CLs based on their dependency on the gene under consideration, in decreasing order. The coordinate on the y-axis indicates the rank position of the gene under consideration from sorting all screened genes based on their fitness scores observed in the CL under consideration, decreasingly. Common-essential genes (RPL8 and RPL22) ranks always among the top fitness scores, resulting in an almost flat trend. The vertical dashed line indicates the 90th percentile of dependency on the gene under consideration. F. Distribution of all genes’ fitness-rank-positions for the CL at their 90th-percentile of least dependent cell lines, i.e. the dashed vertical line in E). The density of these scores is estimated using a Gaussian kernel and the central point of minimum density is identified. Genes whose score falls below this minimum (i.e. to the left of the gray dashed line) are classified as common-essential by FiPer Fixed.

The second and less stringent CoRe method, implemented in four different novel variants, is the Fitness Percentile (FiPer), which identifies CEGs via a pooled (pan-cancer) analysis of data from large-scale CRISPR-Cas9 knock-out screens, performed in cell lines from multiple tissues/cancer-types [31] (**Fig. 1EF**). For each screened cell line, this approach considers the gene rank positions resulting from sorting all screened genes based on their effect on cell viability upon CRISPR-Cas9, i.e. their essentiality, in decreasing order. FiPer then exploits the intuition that CEGs will always rank among the top essential genes for most cell lines, including those for which the fitness reduction is overall less pronounced.

While ADaM takes as input strictly defined binary scores of gene essentiality and it outputs discrete sets of tissue-specific and pan-cancer CFGs, FiPer takes in input quantitative descriptors of gene essentiality and it outputs a unique set of CEGs, also providing a visual means for quickly assessing the tendency of individual genes to be a CEG.

### The Adaptive Daisy Model

The Adaptive Daisy Model (ADaM) [12] is implemented in the function CoRe.ADaM of CoRe, which takes as input (i) a binary dependency matrix, where rows correspond to genes and columns to samples (screens or cell-lines), with a 1 in position *[i, j]* indicating that the inactivation of the *i*-th gene through CRISPR-Cas9 targeting exerts a significant loss of fitness in the *j*-th sample, i.e. that the *j*-th cell line is dependent on the *i*-th gene; (ii) a reference set of prior known CFGs. Binary dependency matrices encompassing data for hundreds of cancer cell lines can be downloaded from Project Score [30] and used with this function by calling CoRe.download_BinaryDepMatrix.

In order to identify CFGs using data from screening *N* cell lines, the Daisy Model introduced in [10] computes a fuzzy intersection of genes that are essential, i.e. fitness genes, in at least *n*\* cell lines, where this number is defined a priori. ADaM generalizes this approach by (i) exploiting the bimodality of the distributions of the number of genes essential in each number of cell lines (**Fig. 1A**), and (ii) adaptively determining an optimal discriminative threshold of minimal number of cell lines *n*\* that should be dependent on a given gene for calling that gene a CFG.

Briefly, for a binary matrix encompassing gene dependency profiles of *n* cell lines across thousands of screened genes, ADaM computes fuzzy intersections of genes *I*_*n*_, for each *n = 1, …, N*. These fuzzy intersections include genes with at least *n* dependent cell lines according to the input matrix. For each tested *n*, ADaM computes the true positive rate *TPR(n)* yielded by each *I*_*n*_ using the reference CFGs provided in input as positive controls. In parallel, ADaM also computes the number of genes that are expected to be essential in at least *n* cell lines by chance, via random permutations of the input matrix (**Fig. 1B**). Finally, ADaM determines the optimal *n*\* as the largest value providing the trade-off between *TPR(n)* (inversely proportional to *n*) and the deviance of the number of genes with *n* dependent cell lines (directly proportional to *n*) from its expectation (**Fig. 1C**). The genes in the corresponding fuzzy intersection *I*_*n*\*_ are predicted to be CFGs for the cell lines in the input dependency matrix.

As the distribution of genes that are CFGs in a specific number of tissue-lineage/cancer-types is also bimodal [12], this procedure can be executed in a two-step approach on large datasets of cancer dependency profiles, accounting for hundreds of cancer cell lines from multiple tissues, to predict pan-cancer CFGs (**Fig. 1D**). In the first step ADaM predicts tissue-lineage/cancer-type specific CFGs, then it iterates by adaptively determining the minimum number *t*\* of tissue-lineages/cancer-types for which a gene should have been predicted as a specific CFG to be now predicted as a pan-cancer CFG. *t*\* is determined by applying the same algorithm and criteria used to determine the *n*\* across the tissue-lineages/cancer-types specific executions of ADaM (**Fig. 1D**). Particularly, this last operation is performed on a binary membership matrix with genes on the rows, tissue-lineages/cancer-types on the column and a 1 in position *[i, j]* indicating that the *i*-th gene is a CFG for *j*-th tissue-lineage/cancer-type.

All the functions called by CoRe.ADaM are exported and fully documented in the CoRe package. In addition, CoRe is equipped with the CoRe.PanCancer_ADaM wrapper function, implementing the two-step procedure to identify pan-cancer CF genes, and the CoRe.CS_ADaM function executing ADaM on a user-defined tissue-lineage/cancer-type, which can be used on dependency matrices from Project Score [30] and cell line annotations from the Cell Model Passports [35].

### The Fitness Percentile Method

The Fitness Percentile (FiPer) method works in an unsupervised manner. It identifies a set of common-essential genes (CEGs) by executing a single pooled analysis of data from multiple CRISPR-Cas9 screens. In addition, it takes in input a dependency matrix with quantitative fitness effect indicators of screened genes across cell lines.

We have designed and implemented in CoRe four novel variants of this method, all sharing the same initial step, which is executed for each individual gene in the input dependency matrix, in turn. In this step (i) all cell lines are sorted according to their dependency on the gene under consideration in decreasing order; (ii) the rank position of the gene under consideration resulting from sorting all screened genes according to their fitness effect is determined, for each screened cell line; (iii) a curve of the rank positions computed in (ii) is assembled considering the cell lines ordered as in (i): the fitness rank versus dependency percentile curve (FiPer curve, **Fig. 1E**).

It is reasonable to assume that genes involved in fundamental cellular processes (likely to be CEGs, such as RPL8 and RPL22 in **Fig. 1E**) will generally tend to rank amongst the most significant fitness genes for all the screened cell lines, including those that are the least dependent on them. This tendency can be extrapolated from the FiPer curves (thus measured in data coming from multiple CRISPR-Cas9 screens) and used to estimate the likelihood of a gene to be a CEG.

The CoRe.FiPer function implements four different methods to assess this tendency assigning a FiPer score to each gene differently. This is followed by a procedure that finally partitions all screened genes into two groups, with the first one containing the predicted CEGs.

The first method, the *Fixed* percentile (**Fig. 1EF**), considers as the FiPer score of a gene its fitness rank position in the cell line falling at the highest boundary of a very large dependency percentile of cell lines (90th by default). The *Average* method considers the average gene rank position in all the cell lines falling over a very large dependency percentile (90th by default). The *Slope* method fits a linear model onto each gene’s FiPer curve, then considers the slope of such a model as the gene FiPer score. In the final *AUC* method, the FiPer score of a gene is computed as the area under its FiPer curve.

Finally, a density function fitted onto the gene FiPer scores’ observed distribution (which is typically bimodal) using a kernel estimator and the score corresponding to the point of central local minimal density is used as a discriminative threshold to predict CEGs, which will be those with a FiPer score less than or equal to it (**Fig. 1F**).

CoRe includes also the CoRe.VisCFness function which visualises the tendency of a given gene to be a CEG within a dependency dataset provided in input and compares this tendency against that of a positive (RPL8 by default) and a negative (MAP2K1 by default) control, and producing the plots shown in **Fig. 1E**.

### Comparison with existing methods and state-of-the-art sets of core-fitness genes

We compared the sets of CFGs and CEGs predicted by CoRe (through ADaM and all the FiPer variants) when applied to the largest integrative dataset of cancer dependency assembled to date, accounting for 17,486 genes and 855 cell lines from 30 different tissue-lineages and 43 cancer types (the DepMap dataset, **Fig. 2AB**) [19], with state-of-the-art sets of core-fitness genes derived from recent functional genetic screening datasets [10, 12, 33, 34]. We also included in the comparison the output of a logistic-regression based method, part of the recent CEN-tools software proposed in [33] applied to the DepMap dataset (**Tables 1 and 2**).

**Table 1-.**
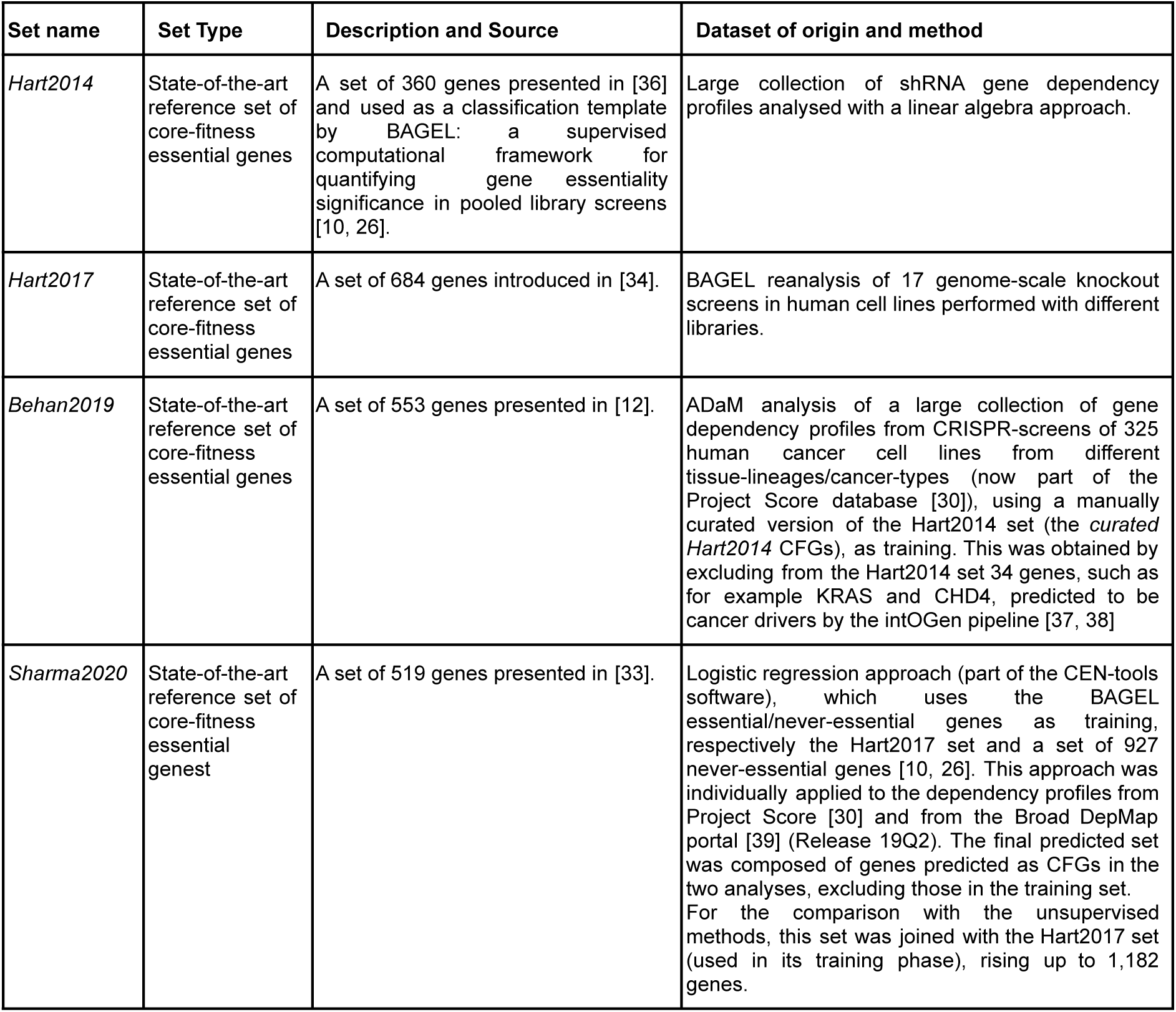
State-of-the-art sets of core-fitness essential genes considered to benchmark CoRe.

**Table 2-.**
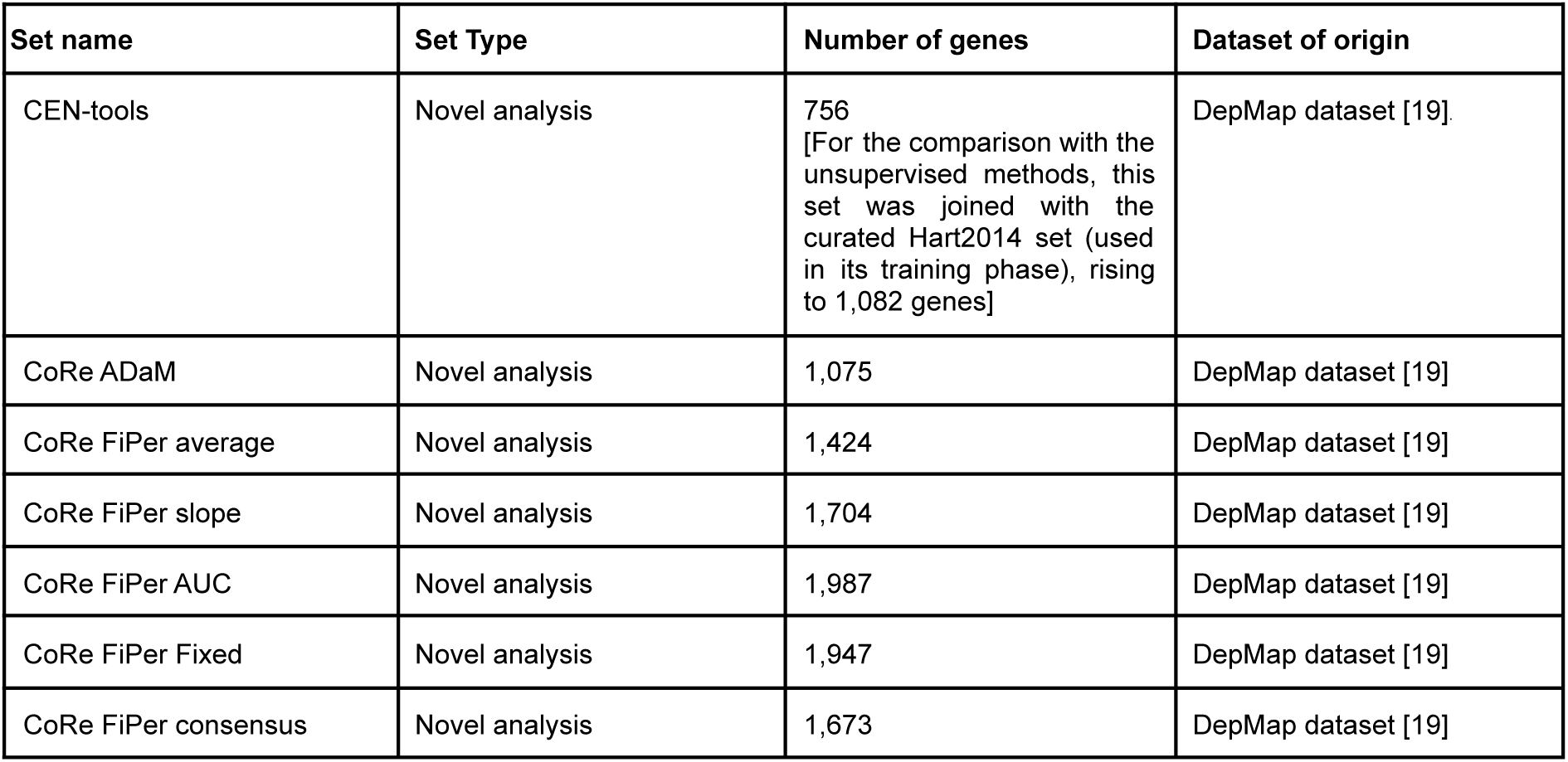
Sets of core-fitness and common-essential genes obtained by novel analyses of the DepMap dataset and considered to benchmark CoRe.

| Set name | Set Type | Number of genes | Dataset of origin |
| --- | --- | --- | --- |
| CEN-tools | Novel analysis | 756<br>[For the comparison with the unsupervised methods, this set was joined with the curated Hart2014 set (used in its training phase), rising to 1,082 genes] | DepMap dataset [19] |
| CoRe ADaM | Novel analysis | 1,075 | DepMap dataset [19] |
| CoRe FiPer average | Novel analysis | 1,424 | DepMap dataset [19] |
| CoRe FiPer slope | Novel analysis | 1,704 | DepMap dataset [19] |
| CoRe FiPer AUC | Novel analysis | 1,987 | DepMap dataset [19] |
| CoRe FiPer Fixed | Novel analysis | 1,947 | DepMap dataset [19] |
| CoRe FiPer consensus | Novel analysis | 1,673 | DepMap dataset [19] |

**Figure 2-.**
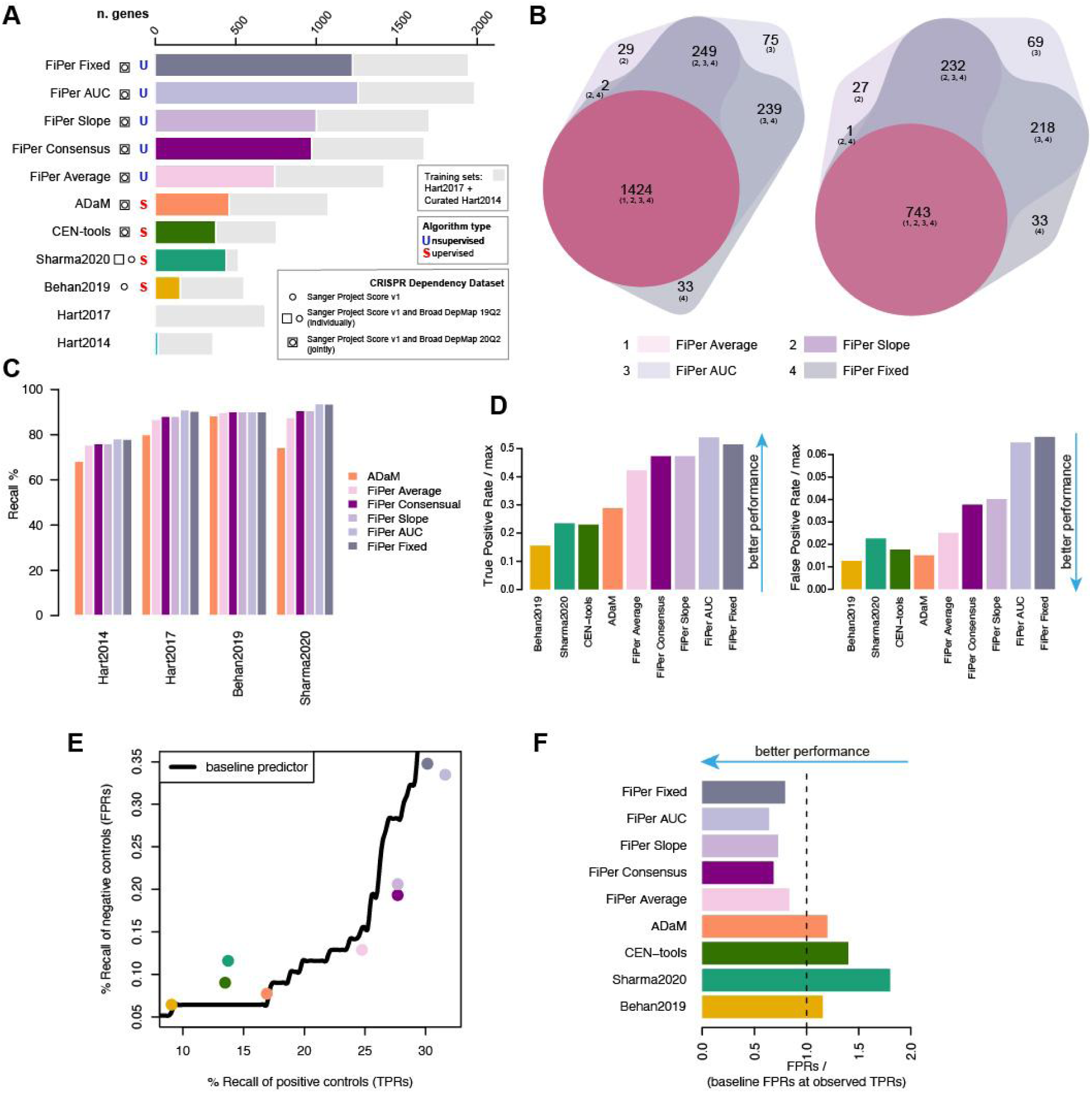
– Comparison of CoRe output with state-of-the-art sets and methods. **A**. For each method predicting core-fitness essential genes (CFGs), common-essential genes (CEGs), or state-of-the-art (SOA) sets of CFGs, the overall length of the bar indicates the total number of genes, whereas the length of the coloured bar indicates the total number of predicted genes not included in any of the training sets. Squares/circles indicate the dataset analysed by each method or used to derive the considered SOA set, and letters indicate the nature of the method, i.e. (S)upervised or (U)supervised. **B**. Comparison of common-essential gene sets predicted by the four variants of the FiPer method (left) and considering novel hits only, i.e., excluding any gene belonging to any of the training sets (right). **C**. Recall of SOA sets of CFGs genes across CoRe methods’ predictions. **D**. True and False positive rates (TPRs/FPRs) of independent true/negative controls across SOA sets of CFGs, CoRe and other methods, relative to the maximal TPRs/FPRs attainable by the basal daisy model (DM) predictor of CFGs. **E**. Performance assessment accounting for set sizes. Each point corresponds to a different method or SOA set, with coordinates indicating their TPR/FPR, respectively along x- and y-axis. Black curve indicates the FPRs obtained by a baseline DM predictor at given TPRs. **F**. FPRs of all tested methods and SOA sets of CFGs relative to baseline performances. The length of each bar indicates the ratio between the FPR of the method or set under consideration and that of the baseline DM classifier at a TPR equal to that observed for the method or set under consideration.

For the training phase of CEN-tools, we used the curated Hart2014 CFGs [12] (which we also used as reference set of positives while running ADaM), and the BAGEL never-essential genes [10], also curated as described in [12] (the curated BAGEL non-essential set).

In order to provide a fair benchmark with respect to sets outputted by the unsupervised methods, we also joined the Sharma2020 set, and the CEN-tools set with the reference CFGs used in their respective training phases, i.e., the Hart2017 set and the curated Hart2014 set. All the compared sets of CFGs and CEGs, the curated Hart2014 essential and curated BAGEL non-essential genes are included in **Additional File 1: Table S1**.

Amongst the predicted CFG sets derived from old and new executions of supervised methods, ADaM yielded the largest number of CFGs (460) not included in any of the training sets (curated Hart2014, Hart2017 and BAGEL non-essentials), when applied to the DepMap dataset (**Fig. 2A**). The Sharma2020 set ranked second (with 441), followed by the novel execution of CEN-tools (with 379) (**Fig. 2A**). As expected, all these sets, included more novel CFGs than Behan2019 (157 novel CFGs), likely due to its derivation from a sensibly smaller cancer dependency dataset (325 cell lines against 855 for ADaM and CEN-tools, and 325 + 489 for Sharma2020, **Fig. 2A**).

The 4 variants of the CoRe FiPer method yielded much larger and highly concordant sets of predicted CEGs (median = 1,825.5, min = 1,424 for FiPer average, max = 1,987 for FiPer AUC, **Fig. 2B**), as well as novel hits (median = 1,115, min = 743 for FiPer average, max = 1,262 for FiPer AUC, **Fig. 2B**). The set of CEGs predicted by FiPer average was included in those predicted by all the other FiPer variants. For this reason, we decided to assemble a 5th FiPer set by intersecting the output of FiPer Slope, AUC and Fixed: the FiPer consensus set. This yielded 1,673 genes, of which 975 were novel hits (**Fig. 2A**).

As a first exploratory analysis, we verified that all the sets of CFGs/CEGs outputted by the CoRe methods covered most of the state-of-the-art sets of CFGs (ADaM median Recall across prior known sets: 77.24%, FiPer median Recall across prior known sets, averaged across variants: 89.31%, **Fig. 2C**). Furthermore, while comparing overall CFG/CEG sets similarities, we observed three major clusters composed respectively by (i) the sets outputted by the FiPer variants, then (ii) Sharma2020, CEN-tools (both joined with respective training sets) and ADaM sets, and (iii) Hart2014, Hart2017 and Behan2019 sets (**Additional File 2: Figure S1**). Taken together, these results suggest that the ADaM, CEN-tools and Sharma2020 sets might include similar numbers of novel CFGs, thus potentially extending in a similar way the other state-of-the-art CFG sets.

To investigate and compare true/false positives rates of the putative novel CFG/CEGs, we assembled, respectively, (i) a set of prior known CFGs (not included into any of the training sets) curated in [19, 21] using data from MsigDB [40], to be used as positive controls, and (ii) considered genes not expressed in human cancer cell lines (using data from the Cell Models Passports [35]) or whose essentiality is statistically associated with a molecular feature (thus very likely to be linked to specific molecular contexts) [19] as negative controls (Methods, **Additional File 3: Table S2**). Both these sets are independent from the DepMap dataset.

Of the CGFs outputted by the supervised methods, ADaM had the best TPR, covering 29% of the positive controls screened in the DepMap. Sharma2020 ranked second (23.4%) followed by CEN-tools (23%) and Behan2019 (15%) (**Fig. 2D**). The median TPR for the FiPer variants was 47%, with FiPer AUC ranking first (54%) and FiPer Average last (42%). In terms of FPRs, Behan2019 performed the best, covering only 1.2% of the negative controls included in the DepMap dataset. ADaM ranked second (1.5%), followed by CEN-tools (1.7%) and Sharma2020 (2.3%). The median relative FPR for the FiPer variants was equal to 4% with FiPer average performing best (2.5%) and FiPer fixed worst (7%).

To account for differences in set sizes, which impact the observed TPRs/FPRs, we sought to compare the observed FPRs with those expected when using a baseline daisy model (DM) predictor of CFGs on the DepMap dataset, considering as the DM thresholds *n*\* the *n* providing the observed TPRs of independent positive controls (**Fig. 2E** and **Additional File 4: Fig S2**).

When considering the supervised methods, CoRe outperformed both CEN-tools and Sharma2020, yielding better ratios of FPRs with respect to those obtained at the observed TPRs by the DM (1.1 and 1.2 respectively for Behan2019 and ADaM, against 1.4 for CEN-tools and 1.8 for Sharma2020 (**Fig. 2F**)). Much better performances were obtained by the FiPer variants (median FPR / baseline ratio = 0.72) with FiPer AUC performing the best (0.64) and FiPer average the worst (0.83).

Optimal sets of CFGs/CEGs are expected to be essential in a vast majority of cancer cell lines: they have an average large negative impact on cellular fitness upon inactivation and are constitutively expressed in non-diseased tissues.

To evaluate these properties across the output of compared methods and state-of-the-art sets, we first measured the median number of cell lines dependent on the predicted sets of CFGs/CEGs (**Fig. 3A**). This was generally large for all the supervised methods, with the Behan2019 CFGs being essential (scaled fitness score < −0.5, Methods) in a median percentage of 99.8% cell lines of the DepMap dataset, followed by CEN-tools (98.9%), ADaM (98.1%) and Sharma2020 (96.8%). As expected, the CEGs yielded by the FiPer variants, were generally essential in smaller but still large percentages of cell lines (grand median = 82.3%, min = 70.2% for FiPer AUC - max = 92% for FiPer average). Nevertheless, when looking at the *n*\* thresholds required by the baseline DM to attain the observed TPRs across predicted CFGs/CEGs (**Fig. 3B**), among the supervised methods the ADaM set showed again the best ratio between median number of dependent cell lines versus baseline (1.14, 98.1% against 86%), followed by CEN-tools (1.06, 98.9% against 93%), Sharma2020 (1.05, 96.8% against 92%) and Behan2019 (1.01, 99.8% against 98.6%) (**Fig. 3C**). The FiPer variants CEGs showed a median ratio between number of dependent cell lines versus DM thresholds at same TPR that was generally strikingly large across methods (median = 2.62, max 4.26 for FiPer AUC - min 1.95 for FiPer average).

**Fig. 3-.**
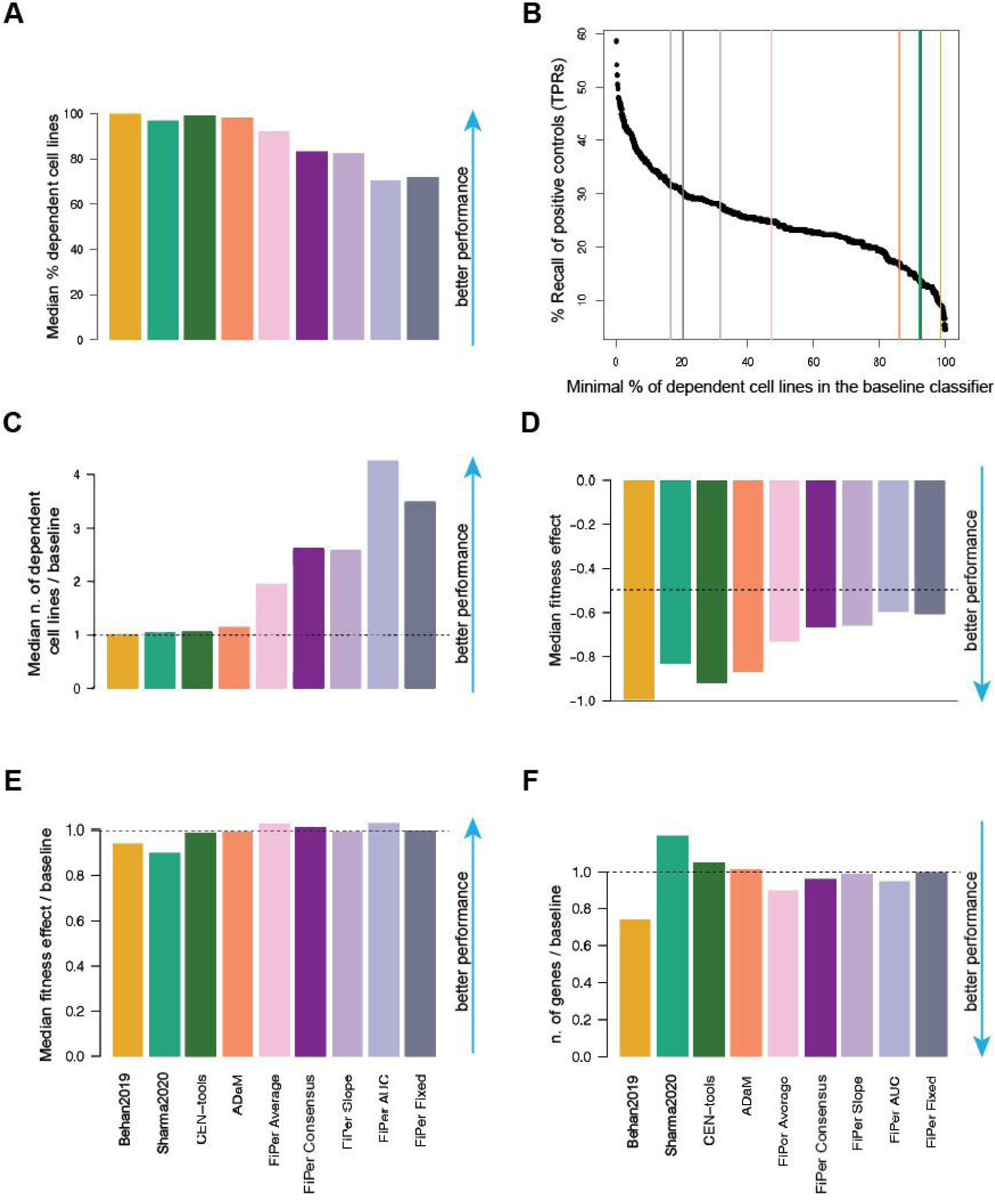
Fitness effects of CFG sets across cell lines. **A**. Median percentage of cell lines in which the genes in the predicted sets or core-fitness gene (CFG) or common-essential gene (CEG) sets are significantly essential. **B**. Threshold of minimal number of dependent cell lines *n* required by the baseline daisy model predictor (DM) to attain the true positive rates (TPRs) observed across tested methods. **C**. Ratios between median numbers of dependent cell lines for predicted sets divided by the threshold *n* of the DM to attain their TPRs. **D**. Median fitness effect exerted by the genes in the predicted CFG/CEG sets. **E**. Ratio between the median fitness effect in D and the median fitness effect exerted by the DM at the observed TPRs. **F**. Ratio between the number of genes in the predicted sets and those predicted by the DM at the observed TPRs.

The proximity to 1 of all the ratios for the supervised methods indicate that they all implicitly discover the DM’s optimal *n*\*. ADaM goes further and selects a set of genes providing a TPR that would require a much lax minimal number of dependent cell lines to be achieved by the DM, thus resulting in an increased FPR. Furthermore, in these circumstances, the unsupervised methods massively outperform the supervised ones, showing the effectiveness of the FiPer criteria used to pick CEGs.

Next, we measured the median scaled fitness effect of the predicted CFGs/CEGs across cell lines, and we find it comfortably below −0.8 -- i.e. 80% of the median effect for curated Hart2014 (Methods) -- for all the supervised methods (strongest effect = −0.99 for Behan2019, weakest for Sharma2020 = −0.83) and below −0.5 -- i.e. half the fitness effect of the curated Hart2014 -- for the FiPer variants (strongest for FiPer average = −0.73, weakest for FiPer AUC= −0.59) (**Fig. 3D**).

Nevertheless, when comparing these values with their equivalent for the CFGs predicted by the baseline DM at the observed TPRs (excluding genes belonging to the training sets), ADaM was again the best performing supervised method (ratio between median fitness effect and baseline = 0.99), followed by CEN-tools (0.98), Behan2019 (0.93), and Sharma2020 (0.89). The median ratio for the FiPer variants was equal to 1.01 with FiPer AUC performing best (1.02) (**Fig. 3E**).

Finally, we found that all the compared methods predicted sets of CFGs/CEGs that were constitutively expressed in normal tissues at similar median levels (**Additional File 5: Fig. S3**). In addition, the CFG sets’ cardinality was systematically comparable or lower than that of CFG sets outputted by the baseline DM at the observed TPRs, with the exception of Sharma2020 and CEN-tools (**Fig. 3F**). Thus, these two sets were confirmed to be suboptimal and predicting larger numbers of CFGs with respect to the baseline DM but with worse FPRs at the observed TPRs (**Fig. 2EF**).

All these results were confirmed when the benchmark analyses were extended to the Hart2014 and Hart2017 sets, adding to CEN-tools and Sharma2020 their corresponding positive training sets and not excluding training set genes from positive/negative controls (thus considering 905 positive and 8,040 negative controls - of which respective 466 and 695 are in the DepMap dataset) (**Additional File 6: Figure S4**).

When considering all state-of-the-art sets of CFGs and supervised methods, we observed again that ADaM provides the best TPRs and FPRs (both absolute and relative to baseline, **Fig. 4A-D**).

**Fig. 4-.**
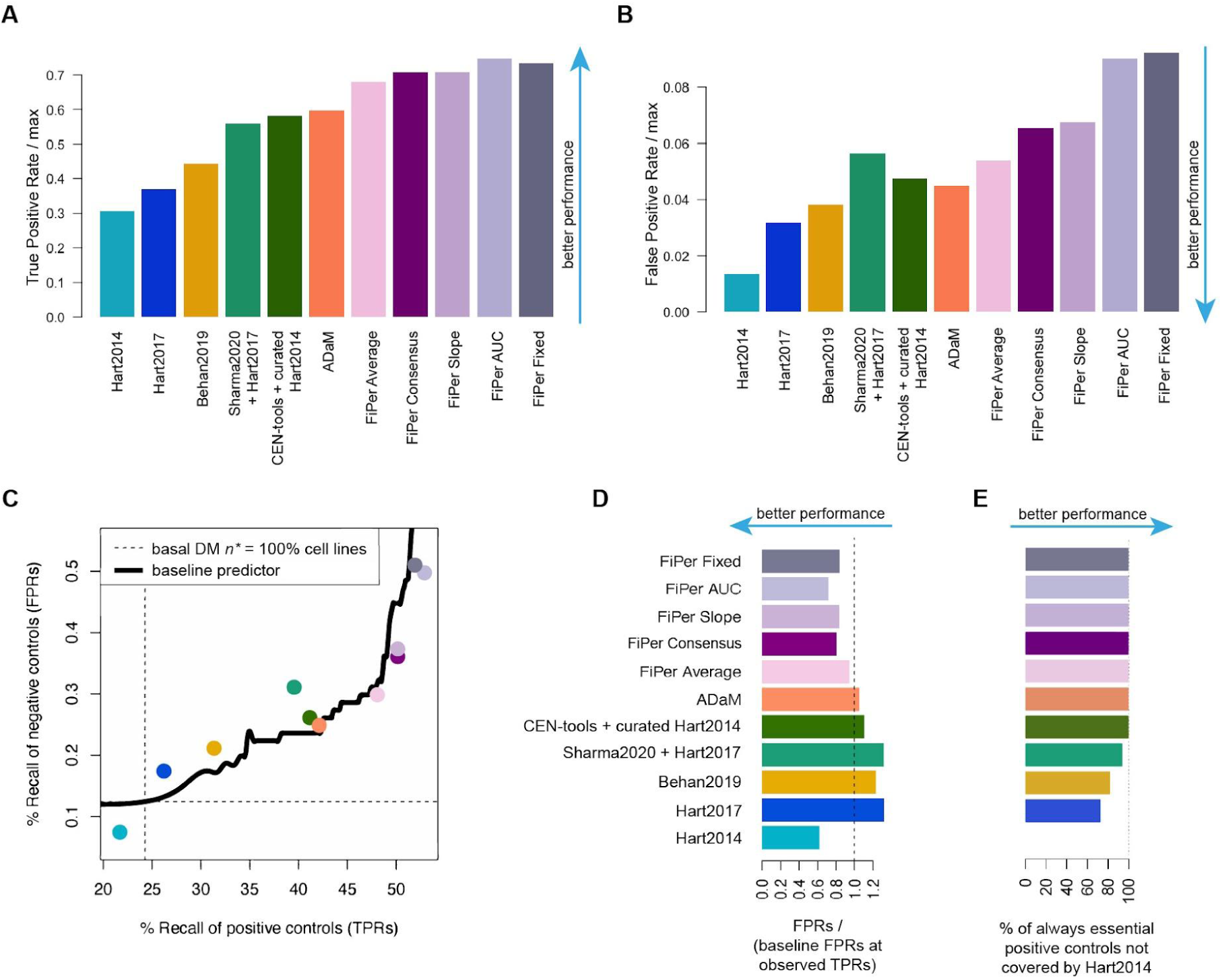
Performances of tested methods when accounting for genes in the training sets. **AB**. True and False positive rates (TPRs, FPRs) of independent true and negative controls across state-of-the-art (SOE) sets of core-fitness essential genes (CFGs), and sets outputted by CoRe and other methods, relative to the maximal TPRs/FPRs attainable by a basal daisy model (DM) predictor of CFGs. **C**. Performance assessment accounting for set size. Each point corresponds to a different method or SOA set, with coordinates indicating their TPR and FPR, respectively, along the x- and y-axis. The black curve indicates the FPRs obtained by a baseline DM predictor at given TPRs. **D**. FPRs of all tested methods and SOA sets of CFGs relative to baseline performances. The length of each bar indicates the ratio between the FPR of the set under consideration and that of the baseline DM classifier at a TPR equal to that observed for that set. **E**. Recall of positive control genes that are essential in 100% of the cell lines in the DepMap dataset and are not covered by the Hart2014 set, across all benchmarked sets.

The Hart2014 set showed the best FPRs versus baseline ratio, although this had to be extrapolated. In fact, this set had a TPR (21.7%) that was lower than that of the baseline DM classifier at the most stringent *n*\* threshold (TPR = 24%, for 343 CFGs that are significantly essential in 100% of the screened cell lines) (**Fig. 4C**), and strikingly did not include 66 positive controls that are significantly essential in all the cell lines of the DepMap dataset (**Fig. 4E**). These 66 genes were all covered by all the methods executed on the DepMap dataset and only partially recalled by the Hart2017 (73%), the Behan2019 (82%) and the Sharma2020 (94%) sets.

Taken together, these results strongly indicate that the CFGs derived from the DepMap dataset reliably extend state-of-the-art CFG sets and that, among those derived with supervised methods, the ADaM set is the most robust one. This was also confirmed in terms of number of cell lines dependent on the predicted CFGs (**Fig. 5AB**) and their median fitness effect (**Fig. 5CD**), relative to baseline performances.

**Fig. 5-.**
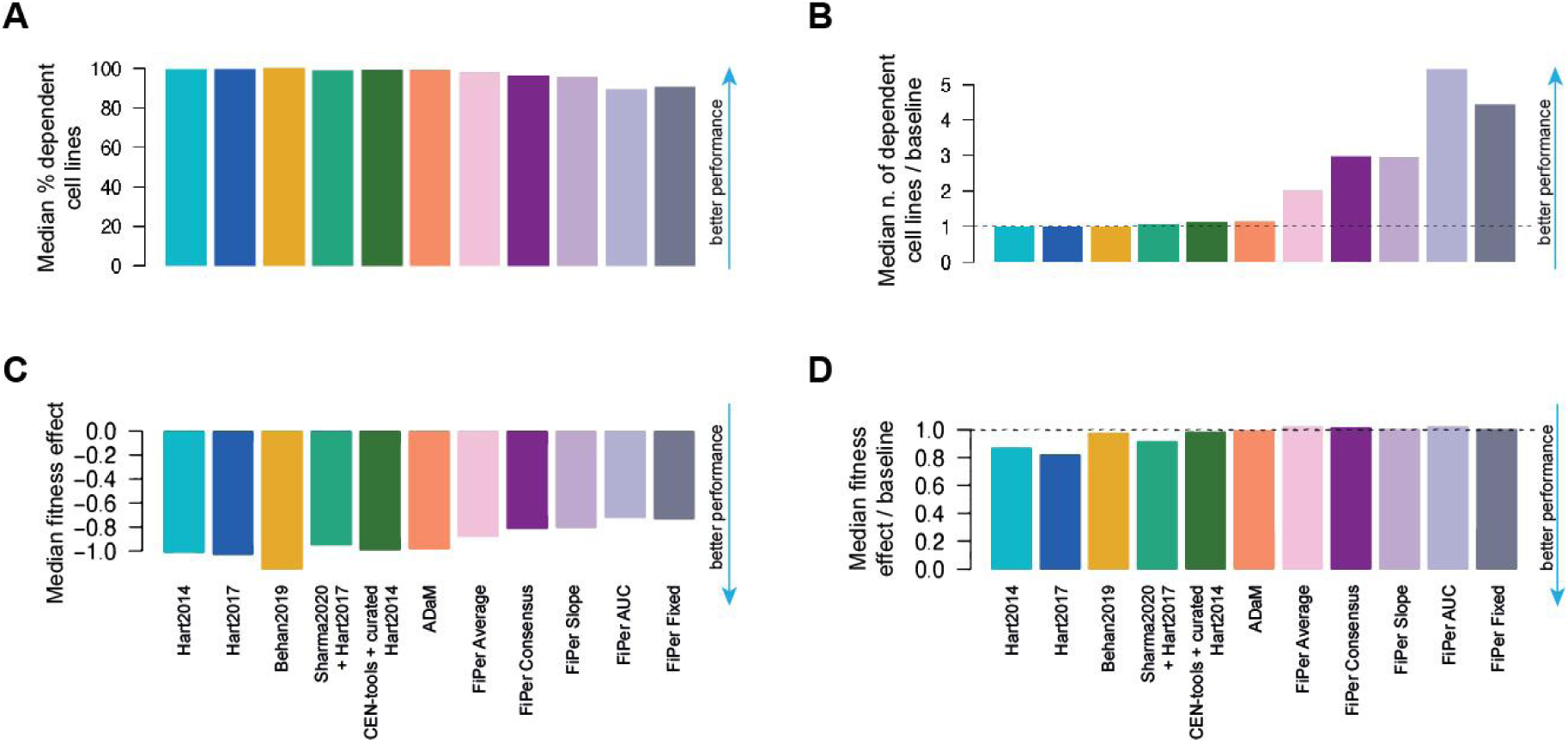
Comparison between CFG/CEG sets’ essentiality profile when accounting for genes in the training sets. **A**. Median percentage of cell lines in which the genes in the predicted sets or core-fitness gene (CFG) or common-essential gene (CEG) sets are significantly essential. **B**. Ratios between median numbers of dependent cell lines for predicted sets divided by the threshold *n* of the baseline daisy model predictor (DM) to attain their TPRs. **C**. Median fitness effect exerted by the genes in the predicted CFG/CEG sets. **D**. Ratio between the median fitness effect in D and the median fitness effect exerted by the DM at the observed TPRs.

### Methods’ performances using an independent cancer dependency dataset

We sought to compare the CGFs and CEGs outputted by the considered methods in terms of their median fitness effect across multiple screened models when using an independent cancer dependency dataset. To accomplish this, we considered an integrated dependency dataset generated by applying the DEMETER2 model to three large-scale RNAi screening datasets, covering 712 unique cancer cell lines [41], pre-processed as specified in the Methods.

Also, in this case, the two versions of the ADaM CFGs sets outperformed the other supervised methods both in terms of absolute grand median fitness effect (−0.79 and −0.61, respectively, for Behan2019 and ADaM, versus −0.6 and −0.5, respectively for CEN-tools and Sharma2020) and ratio with respect to baseline DM (0.98 and 0.96, respectively for ADaM and Behan2019, versus 0.94 and 0.76, respectively for CEN-tools and Sharma2020, **Additional File 7: Figure S5**). As we previously observed, the FiPer variants’ CEGs showed an overall milder grand median fitness effect (median = −0.36) but much better ratios with respect to baseline (median = 0.99).

### Functional characterisation of predicted sets of Core-fitness-essential and Common-essential genes

We performed a systematic statistical enrichment analysis of gene families across all sets of CFGs and CEGs considered in our benchmark, to functionally characterise them. This yielded a set of 13 families significantly enriched (FDR < 5%) consistently across all the state-of-the-art sets of CFGs as well as in the CFGs outputted by all tested supervised methods (**Fig. 6A** and **Additional File 8: Table S3**), thus worthy to be considered as bonafide true positive enrichments in human core-fitness essential genes (the core-fitness families). These families encompass most of the true positive controls used in our benchmark (ribosomal protein genes, proteasome, RNA polymerase [40]), as well as other plausible families, such as proteins involved in the initiation phase of eukaryotic translation [42], chaperonins [43], nucleoporins [44, 45] and less immediate hits, such as AAA-ATPase [46, 47] and WD repeat domain families [48, 49].

**Fig. 6-.**
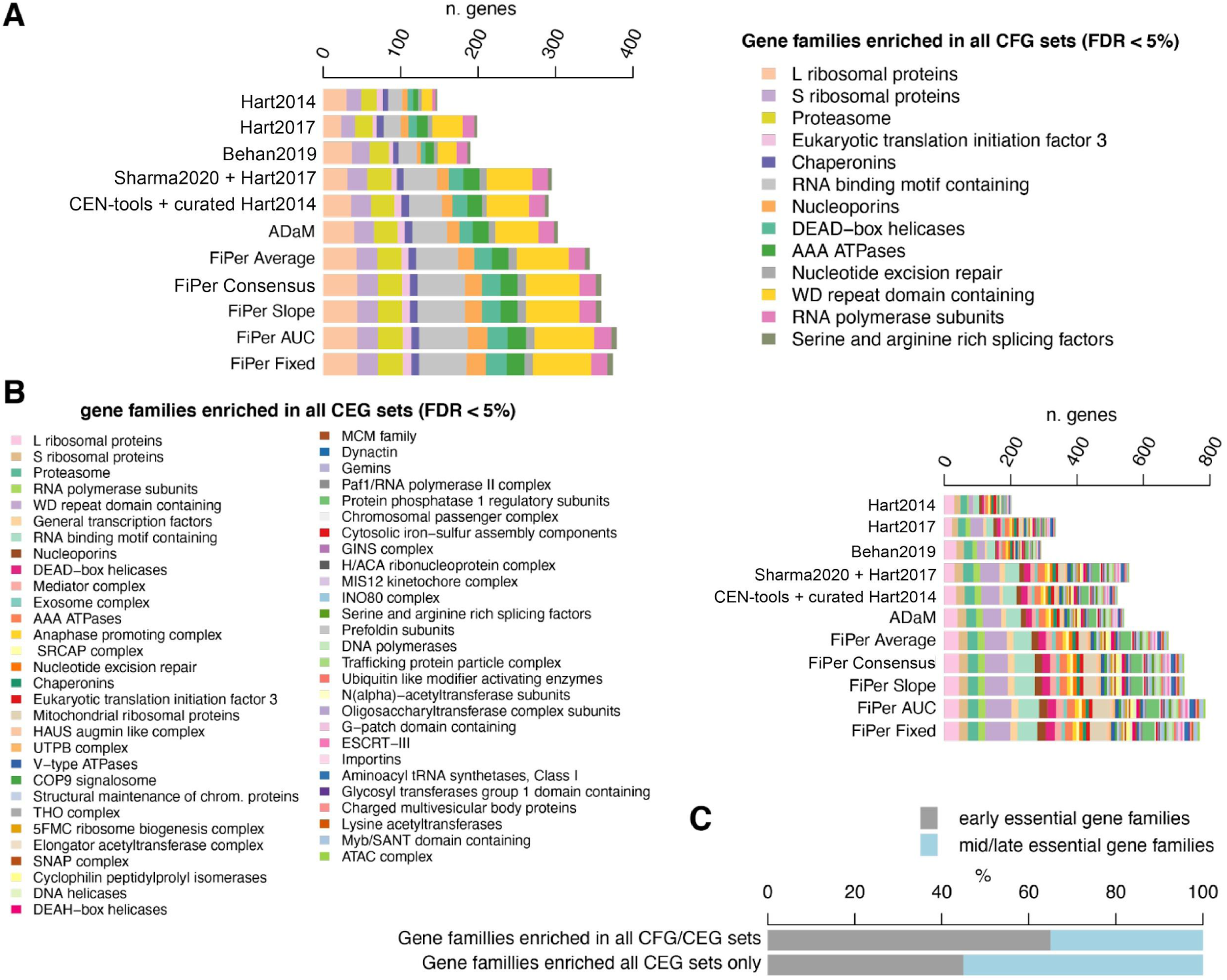
Functional characterisation of predicted core-fitness/common-essential genes. **A**. Gene families consistently significantly enriched (FDR < 5%) across all the state-of-the-art set of core-fitness essential genes (CFGs) and those outputted by the supervised methods. **B**. Gene families consistently and significantly enriched (FDR < 5%) across all the common-essential gene (CEG) sets outputted by the CoRe FiPer variants. **C** Percentage of early and mid/late essential gene families that are also always enriched across CFG and CEG sets or in CEG sets only.

The coverage of these families was much larger for the more recent CFG sets when compared to the state-of-the-art CFGs, with ADaM and Sharma2020 performing best (average Recall across families = 57% and 54%, respectively). The unsupervised methods further extended the coverage of these gene families with average Recalls ranging from 63% (for FiPer average) to 68% (for FiPer AUC), with a median of 65%.

57 gene families were significantly enriched (FDR < 5%) consistently across the CEG sets outputted by the FiPer methods (**Fig. 6B**). These included all the 13 core-fitness families plus 44 additional groups (the common-essential families) such as COP9 signalosome [50, 51], mediator complex [52], SNAP complex [53, 54] and prefoldin subunits [55], to name a few.

When comparing the predicted CFG and CEG sets with the gene-essentiality timing characterisation presented in [56], we observed in the former more genes exerting a negative fitness effect at an early time point upon knock-out (early-essential genes), whereas the latter included more families enriched in genes whose effect on fitness can be detected only at a later time point (late-essential genes) (**Fig. 6C**), such as exosome complex [57], dynactin [58] and ubiquitin-like modifier activating enzymes [59, 60].

### Evaluation of core-fitness gene sets as template predictors of cell line specific essential genes

We performed a final analysis evaluating each state-of-the-art set of core-fitness essential genes (CFGs), and those outputted by CEN-tools and ADaM when applied to the DepMap dataset, as a template classifier of cell line specific essential genes with BAGEL: a widely used bayesian method to estimate gene essentiality significance in pooled CRISPR-cas9 screens [26].

To this aim, we analysed with BAGEL the dependency profiles in the DepMap dataset generated at Sanger, and preprocessed with CRISPRcleanR [21] (Methods), obtaining 7 instances of BAGEL Bayes Factor (BF) matrices, quantifying the likelihood of each gene to be essential in each cell line, using each of the benchmarked set in turn as positive reference set of essential genes in the BAGEL classification template. To evaluate the robustness of the obtained cell line specific BFs we assembled a set of cell line specific positive/negative essential-gene controls.

As positive control, we considered putative oncogenetic dependencies arising from oncogenes (from [38]) found mutated or copy number amplified in a cell line (using data from the Cell Model Passports [35]), whereas wild-type and non-expressed (FPKM < 0.1) oncogenes were considered as negative controls (**Additional File 9: Table S4**).

Then, we assessed the 7 BF matrices, pooling all included values together and considering them as a unique rank-based predictor (the larger the BF the higher the likelihood of a gene to be essential) of cell line specific essential genes, by means of receiver operating characteristic (ROC) analyses (Methods). Particularly, for each benchmarked set we computed the area under the BF-rank induced precision-recall curve (AUPRC) (**Fig. 7A** and **Additional File 10: Figure S6**) and the recall of positive controls at 5% FDR (**Fig. 7B**). All the sets of CFGs outputted by CEN-tools and CoRe applied to the DepMap dataset (**Table 2**) outperformed the state-of-the-art sets of CFGs, showing a better ability in detecting as significantly essential mutated oncogenes, when used as a template for BAGEL. Above all, ADaM achieved the highest recall at 5% FDR (Methods).

**Fig. 7-.**
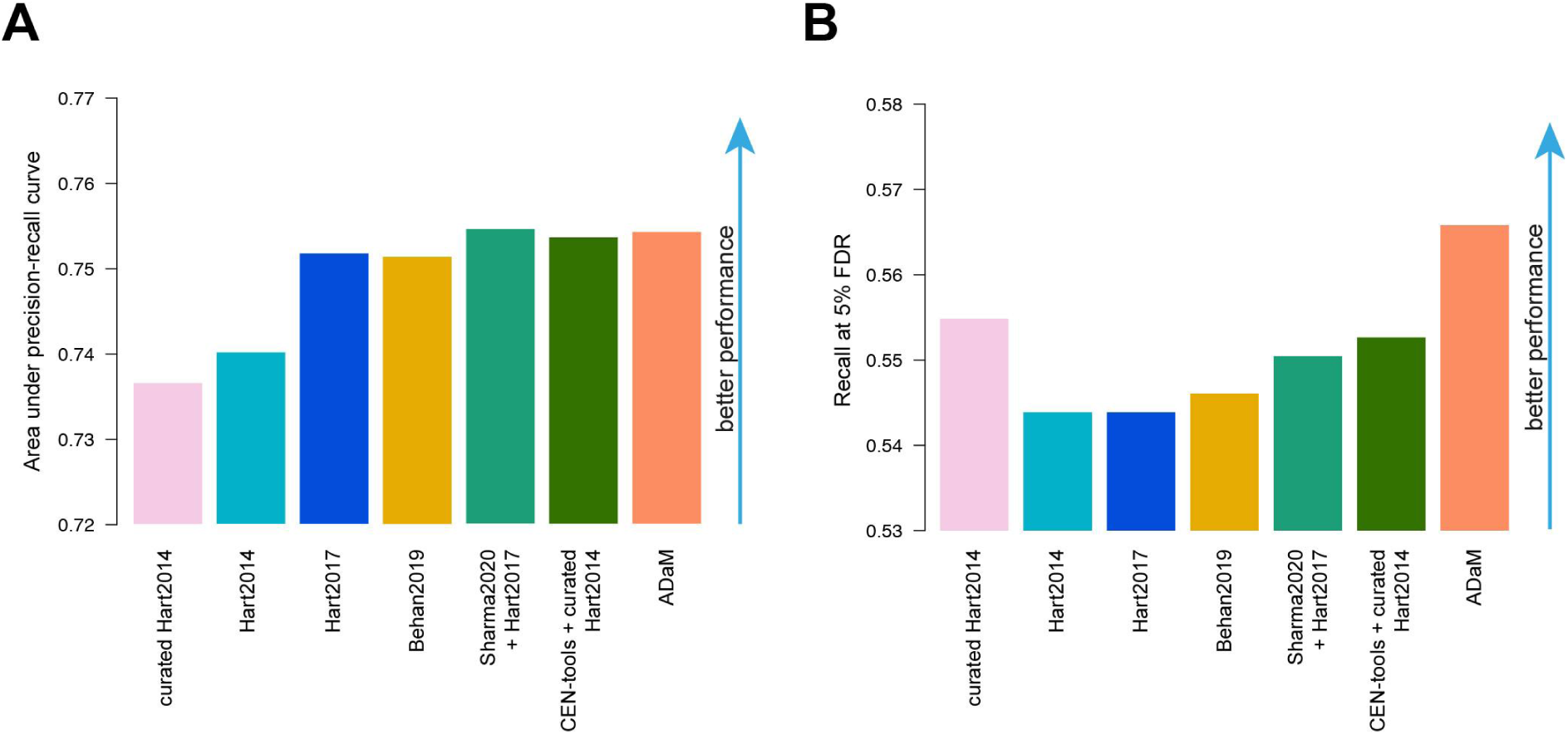
Performances of the benchmarked sets as template classifiers of cell line specific essential genes. **A**. Area under precision-recall curve obtained when predicting cell line specific oncogenetic addictions versus not expressed oncogenes with rank-based classifiers yielded by gene essentiality Bayesian factors. These are computed by BAGEL using each of the benchmarked sets as positive classification template. **B**. Recall of cell line specific oncogenetic addictions at 5% FDR of not expressed oncogenes yielded by each benchmarked set when used as for A.

### Computational efficiency

We measured and compared running times of the benchmarked methods applied to the DepMap dataset, on different operating systems as well as on Google CoLab, a Jupyter notebook service hosted by Google servers (**Table 3**). The CoRe FiPer methods were between 16 (FiPer slope vs ADaM on Ubuntu 16.04 LTS) to 98 (FiPer fixed vs ADaM on CoLab) times faster than ADaM and between 31 (FiPer slope vs CEN-tools on Ubuntu 16.04 LTS) to 123 times (FiPer fixed vs CEN-tools on CoLab) faster than CEN-tools. Across FiPer variants, the slope one was the slowest, probably due to fitting of a linear regression model to a discrete distribution of gene fitness-rank-positions. Nevertheless, FiPer’s running time was still significantly lower than ADaM and both outperformed CEN-tools, which was the method with the longest running times, invariantly across operating systems.

**Table 3-.**
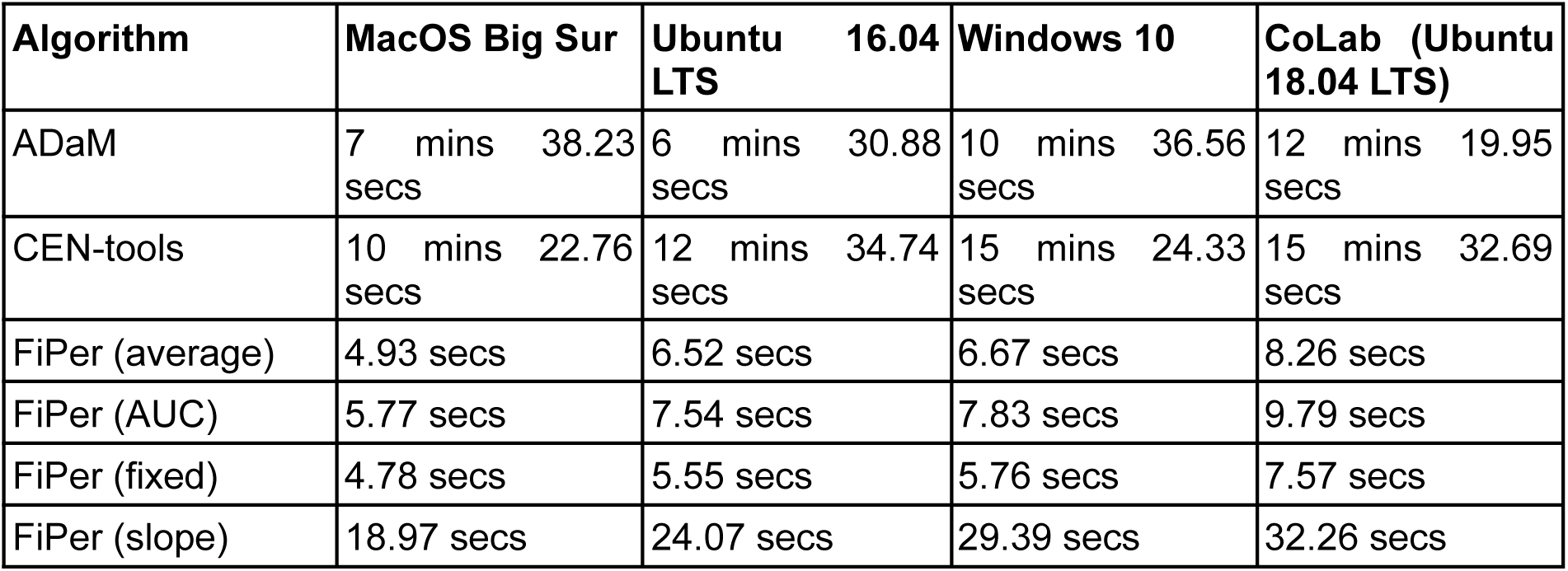
Computational efficiency across methods. Assessments of running time of the six compared methods when executed on different operating systems and on Google Colab.

## Discussion

We introduced CoRe: an open-source R package implementing both existing and novel methods for the identification of core-fitness essential genes (CFGs) --at two different levels of stringency--from joint analyses of multiple CRISPR-Cas9 pooled recessive screens. We robustly and extensively benchmarked CoRe against state-of-the-art sets of core-fitness genes and other CFGs discovery methods, using the largest integrative dataset of cancer dependency to date. We observed that the sets of core-fitness essential and common-essential genes (CEGs, outputted by the less stringent methods) predicted by CoRe are much more comprehensive and robust, in terms of true and false positive rates (TPRs, FPRs) both absolute and relative to a baseline classifier. For the latter, we considered a simple baseline daisy model (DM) model [10] outputting as CFGs those genes exerting a negative effect on fitness upon CRISPR-cas9 targeting in at least an optimal minimal number of screened models, which is known a priori. We also demonstrated that both CoRe and other methods can implicitly detect this optimal DM threshold, with the CoRe methods going much further and accurately predicting sets of genes that are essential in numbers of cell lines that are larger than this threshold. This is much more evident for the less stringent methods implemented in CoRe (i.e., the FiPer variants), thus showing the effectiveness of their underlying algorithm (based on genes’ fitness percentile curves), which selectively picks likely true CEGs. Particularly, across these variants, the FiPer AUC method performs the best even when compared to a consensus set of CEGs obtained by intersecting the output of all the other FiPer variants. Consistently, AUC is the FiPer variant implemented/executed by default by CoRe. However, the other variants are also implemented in CoRe and can be executed for reproducibility purposes.

Contrary to other methods, the sets of CFG/CEG predicted by CoRe are also smaller than those outputted by a baseline DM predictor attaining the same true positive rates, and our benchmark results were all confirmed when extending the analysis to gene sets used in the training phase of at least one of the compared methods, and when considering an independent RNAi based cancer dependency dataset.

Furthermore, we found that the CoRe CFGs/CEGs extend gene families covered by previous state-of-the art sets and methods, with the FiPer methods being able to detect more subtle yet consistent fitness effects and core late essential genes. Finally, the CoRe CFGs/CEGs are all constitutively expressed in non-diseased tissue, pointing to the primary role which these genes play inside the cell. Indeed, it has been shown that higher essentiality is correlated with higher expression and association in important biological pathways [61].

Importantly, our final benchmark analysis also suggests that the CFGs yielded by our novel analyses of the DepMap dataset might be better suited than the reference positive control sets currently used [34, 36] as positive predictor template when estimating cell line specific essential genes with a supervised classification method, such as BAGEL [26].

The development of new tools exploiting the wealth of data currently being generated from CRISPR screens is of paramount importance [62]. Paired with the generation of new data from large efforts and collaborative endeavours, such as for example the Cancer Dependency Map [32, 63], this will be vital for identifying new oncology therapeutic targets, as well as for the characterisation of novel human core essential genes. Nevertheless, another key need is to couple CRISPR screening data with other genetic and molecular information of the screened models and data from ‘normal’ samples. A major reason for this is that a context-specific essential gene in a given cancer genetic background might be, for example, too toxic if suppressed *in vivo* or, in the opposite case, a gene characterized by a pan-essentiality profile in cancer might show reduced on-target toxicities [64].

## Conclusions

The identification of core-fitness genes has important implications in different areas of the life sciences: from drug discovery and cancer therapy to the study of genetic networks. However, different strategies are required according to the type of biological question being investigated. From this perspective, the utility of CoRe is twofold. In fact, when performing functional genetic studies or aiming at identifying novel CFGs, we recommend adopting a more stringent approach, such as ADaM, which can guarantee higher confidence. On the other hand, when the focus is on the identification of new therapeutic targets, thus, to seek new promising context-specific essential genes, the opposite is true. Therefore, applying a less stringent algorithm, such as the FiPer method (particularly the FiPer AUC) allows a larger number of genes to be classified as common-essentials, thus ruling out confounding genes that may skew the outcome of the analysis.

In addition, the CoRe workflow can be adapted to users’ needs and contingencies and it is compatible with many pre-processing methods and tools to estimate fitness effect significance. For example, the recently introduced Chronos tool [65] (accounting for cell population dynamics while estimating gene essentiality) could be used instead of CERES [66]. In addition, when copy number alteration profiles are not available for the screened models, the unsupervised method CRISPRcleanR [21] could be used to correct for gene-independent responses to CRISPR-Cas9 targeting. Furthermore the recent BAGEL2 tool [67] can be used in the initial binarization of essentiality scores, required for ADaM.

Finally, where sufficient data is available, i.e. enough screened models, the algorithms implemented in CoRe could be used to analyze specific subsets of cancer cell lines hosting certain molecular features (e.g. KRAS mutations in colorectal carcinoma), allowing identifying/comparing subtype specific core-fitness genes, which would be of particular interest for translational cancer research.

With the increasing availability of comprehensive cancer dependency maps [32], tools such CoRe will be arguably more and more needed in the future, and they will contribute translating data and findings from such efforts into novel therapeutic target candidates.

## Methods

### Source of reference essential and non-essential genes

The Hart2014 (used as reference sets of prior known essential genes) and the BAGEL non-essential set (used as reference sets of prior known non-essential genes) are introduced in [36] and are derived from a collection of shRNA screens across 72 breast, ovarian and pancreatic human cancer cell lines [68]. We have further curated these sets by removing high confidence cancer driver genes as described in [12]. The Hart2017 (used to train the logistic regression model in [33] together with the BAGEL non-essential set) set was derived in [34] from a reanalysis of 17 genome-wide CRISPR-Cas9 knock-out employing three different CRISPR libraries [4, 69–71]. All these sets have been independently generated and are independent from the DepMap datasets described below.

### DepMap dataset acquisition and pre-processing

We downloaded the latest version of the integrated Sanger and Broad essentiality matrix processed with CERES [66] from the DepMap portal (‘https://www.depmap.org/broad-sanger/integrated_Sanger_Broad_essentiality_matrices_20201201.zip‘). Among the 908 cell lines/columns, 51 were found to contain missing values and were thus removed. We then kept only the cell lines with an associated cancer tissue in the Cell Model Passport (annotation file version 20210326, https://cog.sanger.ac.uk/cmp/download/model_list_20210326.csv.gz), totalling to 855. This step is required to run ADaM tissue-wise. The dataset was then scaled column-wise to have the median of curated BAGEL essential gene fitness scores equal to −1 and the median of curated BAGEL never-essential equal to 0 across all cell lines.

For the execution of ADaM, we binarised the pre-processed CERES dataset considering as essential all genes having a fitness score less than −0.5 in each cell line, otherwise they were considered as non-essential. In addition to directly binarising the fitness score matrix, other strategies might be adopted: for example, use of BAGEL2 [67] for the estimation of gene-level bayesian factors, followed by per-sample scaling by subtracting the 5% FDR threshold between the fitness score distributions of reference sets of essential and non-essential genes, and binarization by assigning 1 to all positive scores, 0 otherwise, as illustrated in [12].

### CEN-tools Logistic Regression execution

We downloaded the CEN-tools package [33] from https://gitlab.ebi.ac.uk/petsalakilab/centools/-/tree/master/CEN-tools. In order to decrease the memory burden for the GitHub repository of the CoRe package, we removed all the python modules and data objects that were not directly called by the LR.py and clustering.R functions, respectively the python script implementing the logistic regression model and the R script performing the subsequent cluster analysis.

In addition, we added a few lines of code to the LR.py script to make it runnable from the command line and compute data objects on the fly. Particularly, CEN-tools uses a python dictionary in pickle format to specify which genes belong to the true positive set (i.e., curated BAGEL essential) or true negative set (i.e., curated BAGEL never-essential). Both scripts were seeded to guarantee reproducibility. All changes applied to the CEN-tools script are detailed in **Additional File 11 - Additional Documentation**.

For the execution of the logistic regression model implemented by CEN-tools we used the curated BAGEL essential and curated never-essential genes for the training phase [12]. Based on the logistic regression, CEN-tools computes a matrix of continuous probability distributions for each gene being essential across cell lines and discretizes them according to the number of bins specified by the user. We adopted 20 bins as this was the default parameter used in the original CEN-tools run [33]. Following the pipeline, the matrix was normalised and genes not included in the training sets were then clustered through k-means using the Hartigan-Wong algorithm [72] around four centres. The silhouette method identified four as the optimal number of clusters according to their probability essentiality profiles: core essential, context-specific, rare-context-specific, and never-essential. The core essential genes are characterized by the highest value of silhouette width and were then used for the downstream benchmarking.

### Execution of ADaM

ADaM takes as input a binarised matrix of fitness essentiality scores. For the identification of tissue CFGs, only the *N* cell lines that are part of the same cancer tissue/type *T* are selected. ADaM then implements a fuzzy intersection *I*_*n*_ composed of genes exerting a significant depletion in at least n cells out of *N*. The threshold *n*\* is obtained in a semi-supervised manner: for each possible fuzzy intersection *I*_*n*_ from *n = 1 to N*, a true positive rate (*TPR(n)*) is computed considering a set *E* of a priori known essential genes as true positives, while *G* is the whole set of screened genes:

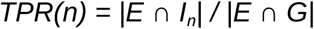

In addition, ADaM computes the log_10_ odd ratio between the observed *I*_*n*_ and its expectation value E(I_n_):

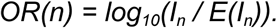

*E(I*_*n*_) is estimated by shuffling the binary matrix column-wise 1000 times. In this way the number of essential genes for every cell line in *T* is preserved. Then *E(I*_*n*_) is defined as the average value of *I*_*n*_ cardinality:

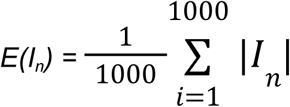

The threshold *n*\* corresponds to the minimal number of cell lines *n* whose *I*_*n*_ provides the trade-off between the two monotonic functions, *TPR(n)* being inversely proportional to *n* and *OR(n)* being directly proportional to *n*.

This is implemented by the wrapper function CoRe.CS_ADaM that subsets the dataset by taking only the cell lines included in the cancer tissue/type of interest using the Cell Model Passport [35] annotation file. We used the annotation file version 20210326.

ADaM was executed using the CERES binarised dataset and using the curated BAGEL essential genes as reference true positives. We set the number of random trials for the generation of the null model to 1000 and ran the algorithm only on those cancer tissues with at least 15 cell lines available as detailed in [12].

### Execution of FiPer variants

We used the function CoRe.FiPer of CoRe. Initially, the method computes a gene-wise cell line ranking *R*_*CL*_, where for each gene g it ranks every cell line cl according to the fitness essentiality score of *g* in *cl*. It also computes a cell-wise gene ranking *R*_*G*_, where for each cell line it ranks every gene according to the fitness essentiality of *g* in *cl*. Then the package implements four different variants of the fitness percentile method:

- Fixed = a distribution of gene fitness-rank-positions in their most dependent n^th^ (determined by the percentile parameter, default is 90^th^) percentile cell line is used in the subsequent step.
- Average = a distribution of average gene fitness-rank-positions across cell lines at or over the n^th^ percentile of most dependent (determined by the percentile parameter, default is 90^th^) cell lines is used in the subsequent step.
- Slope = for each gene g, a linear model is fit on the sequence of gene fitness-rank-positions across all cell lines sorted according to their dependency on g, then a distribution of models’ slopes is used in the subsequent step.
- AUC = for each gene g, the area under the curve (AUC) resulting from considering the sequence of gene fitness-rank-positions across all cell lines sorted according to their dependency on g is used in the subsequent step.

Each FiPer variant outputs a discrete distribution of gene fitness-rank-positions. A gaussian kernel estimator is applied to compute a continuous distribution. The kernel density estimator uses a default bandwidth defined as 0.9 times the minimum of the standard deviation and the interquartile range divided by 1.34 times the sample size to the negative one-fifth power. The distribution is bimodal, and the rank threshold corresponds to the local minimum. All genes having a fitness-rank-position lower than the threshold are classified as CEGs. In the benchmarking, we assessed all the four variants on the full pre-processed CERES matrix to identify the pan-cancer CEGs.

### Benchmark of pan-cancer core-fitness gene sets

We designed a baseline predictor to assess the sets of pan-cancer core-fitness genes computed by each method. This was defined by considering core-fitness those genes essential in at least *n* cell lines for all possible *n*.

Next we assemble sets of priori known positive/negative control essential genes. For the positive controls we collected signatures of genes involved in fundamental biological processes and universally essential genes: such as genes coding for ribosomal proteins, RNA polymerases, histones, or genes involved in DNA replications, etc. curated in [21] and [19] from MsigDB [40]. As negative controls we assembled a set of genes never expressed (fragments per kilobase of transcript per million mapped reads (FPKM) < 0.1) in more than 1,000 human cancer cell lines (from the Cell Model Passports [35]), or whose fitness signal across hundreds of cell lines has a t-skewed normal distribution (according to the normLRT score introduced and applied to an independent shRNA-based cancer dependency dataset in [73]) and it is statistically associated with a genomic marker [19]. Excluding genes included in at least one of the training sets yielded a final set of 408 positive controls and 7,767 negative controls. Of these, 265 positive controls and 555 negative controls were included in the DepMap dataset.

The metrics derived from the baseline predictor were used to assess each CF set. We considered both novel hits, namely the CF sets stripped out of the BAGEL genes used in the training phase of the two CEN-tools runs (i.e. the Hart2017 set, the BAGEL non-essential genes [36], curated BAGEL essential and never-essential genes [12]), as well as in their entirety. The recall of each set was normalised by the maximum recall achieved by the baseline predictor and so was done for the false positive rate. The two coordinates associated with each set were used to perform a cubic spline interpolation [74] and evaluate the balance between the normalised recall and false positive ratios according to the size of the set.

Next, we computed the thresholds required by the baseline predictor to attain the recalls observed by all tested methods and corresponding false positive rates.

### Characterisation of novel pan-cancer core-fitness sets

To identify biologically grounded novel pan-cancer CF sets, we considered the gene families found enriched across all the predicted sets. For every set, we performed a hypergeometric test for each gene family with at least one gene in the CFG set of interest, following the formula:

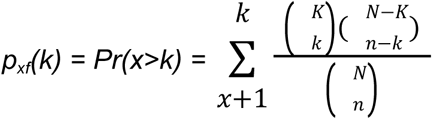

where *p* is the associated probability value of having more genes than observed *k* for a given family *f* in the CFG set under consideration, *K* is the total number of genes in the CFG set associated to any functional family, *N* is the total number of screened genes in the DepMap dataset preprocessed matrix associated to any functional family, *n* is the total number of genes belonging to f and found either in the CFG set or the remaining screened genes.

The *p*-values were then pooled and corrected set-wise using the Benjamini-Hochberg procedure. Gene families with an adjusted *p*-value < 0.05 in each CFG set were deemed significant. The significantly enriched families in common across the supervised methods (i.e., ADaM and CEN-tools in both instances, including also the Hart2014 and Hart2017 sets) were classified as always enriched and the pooled CFG sets used as ground truth. Particularly, we computed the exclusive CEGs in the FiPer AUC set belonging to the always enriched families that were not found in the ground truth. These CEGs were classified as novel hits.

In addition, we repeated the analysis assembling the significantly enriched families in common across the unsupervised methods (i.e. the four variants of the fitness percentile method plus the FiPer consensus set) and showed that unsupervised methods have higher sensitivity in identifying families derived from late time-point essential gene sets [56].

### Benchmark using an independent cancer dependency dataset

We downloaded DEMETER v6 04/20 (available at https://ndownloader.figshare.com/files/11489669), cancer dependency data derived from genome-wide RNAi screens [41]. This dataset was scaled column-wise in order to have the median of curated BAGEL essential gene fitness scores equal to −1 and the median of curated BAGEL never-essential equal to 0 across all cell lines.

For each CFG/CEG set, the median DEMETER fitness scores for every gene across cell lines were derived. In addition, we also derived the median DEMETER fitness scores of the genes included in the CFG sets predicted by the baseline classifier, at the observed TPRs, and computed the normalised scores across sets.

### Retrieval of oncogenetic addictions as bayesian factor templates

We ran BAGEL v115 on the Sanger release 1 cancer dependency dataset (downloaded from: https://score.depmap.sanger.ac.uk/downloads) processed with CRISPRcleanR [21] (shown in [19] to better preserve context-specific essentialities than CERES). As a positive training gene set we used each of the sets among state-of-the-art sets, or CFG sets derived from the supervised methods, in turn, whereas as a negative training gene set we used the curated BAGEL never-essential genes. This led to seven different templates of Bayesian factor (BF) matrices. Each template was scaled cell-wise by subtracting to each gene the 5% false discovery rate (FDR) threshold computed between the BF scores of the two training distributions, for comparability.

Next, we defined a set of cell line specific true positives and negatives to assess the ability of the templates in recapitulating oncogene addictions. First, we assembled a binary matrix summarizing the status of pan-cancer Cancer Functional Events (CFEs) across Sanger cell lines, namely somatic mutations, copy number alterations, and hypermethylation. The binary matrix was then subset to include only genes unambiguously classified as oncogenes in the catalog of driver genes release 2020.02.01 from the IntOGen database. In addition, copy number gains in genomic segments containing ERBB2 or EGFR or KRAS or MYC or MYCN (typically copy number amplified oncogenes in different cancer types), were considered as positive events too. Secondly, we assembled an additional binary matrix where we deemed as positive events oncogenes not expressed in a cell line. We considered oncogenes only instead of including all not expressed genes to avoid unbalanced control sets, favouring the negative controls. By combining the two binary matrices, we obtained three classes:

- Positive instances constituted by oncogenes mutated and expressed in a cell line.
- Null instances constituted either by wild-type and expressed or mutated and not expressed oncogenes in the cell line.
- Negative instances constituted by wild-type and not expressed oncogenes in the cell line.

The positive and negative instances were used on the BF score of each template to assess the area under precision-recall curve and the recall at fixed percentages of FDRs.

### Hardware and software used to compare computational performances

All the analyses were performed on a MacOS laptop with a 2.3 GHz Quad-Core Intel Core i7 processor and 16 GB RAM. The operating system was Big Sur v11.2.3 (20D91). The software was executed in the RStudio IDE v1.3.1073 with x86_64-apple-darwin17.0 platform and R programming language v4.0.2, python scripts were executed using python v3.9.1. For all the methods shown in Table 1 below but ADaM, we used the quantitative pre-processed CERES matrix. The matrix consisted of 17,846 genes and 855 cell lines containing gene fitness scores. Instead, ADaM used a binarized version of the matrix as explained in the previous section. The binary matrix consisted of 8,496 genes, considering genes classified as essential in at least one cell line, and 820 cell lines, considering cell lines from a cancer tissue with at least 15 cell lines in total.

In addition, we executed the CoRe methods and CEN-tools on an ASUS laptop with a 1.80 GHz Quad-Core Intel Core i7 processor and 8 GB RAM. The laptop resources were partitioned 50/50 following a dual-boot configuration, presenting on one end Ubuntu 16.04 LTS as operating system, and Windows 10 on the other. The methods were tested on both operating systems.

## Supporting information

Additional Files - S Figures, S table Legends, documentation

## Declarations

### Ethics approval and consent to participate

Not Applicable

### Consent for publication

Not Applicable

### Availability of data and materials

CoRe is publicly available as an open-source platform independent R package at https://github.com/DepMap-Analytics/CoRe (DOI: 10.5281/zenodo.4813061, license: GPL (>=3)). An interactive vignette, with demonstrations and examples is available at https://rpubs.com/AleVin1995/CoRe. The package includes built-in visualisation and benchmarking functions and their related data objects. It also contains interface functions for downloading and processing state-of-the-art cancer dependency datasets from Project Score [30], as well as updated cancer cell line annotations from the Cell Models Passports [35]. Finally, results from benchmarking CoRe against state-of-the-art sets of CFGs and other CFGs identification methods, with corresponding figures, are fully reproducible executing the Jupyter notebook (also compatible with Google CoLab) available at: https://github.com/DepMap-Analytics/CoRe/blob/master/notebooks/CoRe_Benchmarking.ipynb.

All used data are publicly available at the locations indicated in the main text or in the methods or embedded as native data objects in the CoRe R package.

### Competing interests

MJG and FI receive funding from Open Targets, a public-private initiative involving academia and industry. MJG receives funding from GSK, AstraZeneca, and has performed consultancy for Sanofi. MJG is founder of Mosaic Therapeutics. FI performs consultancy for the joint CRUK-AstraZeneca Functional Genomics Center.

### Authors’ contributions

AV conceived the study, designed, and performed the benchmark analyses, wrote and documented the CoRe package, assembled the interactive vignette and the jupyter notebook, wrote and revised the manuscript. EK wrote and documented the first version of CoRe and revised the manuscript. CP contributed to package writing and documentation and revised the manuscript. UP and RRDL contributed to the design of the benchmark analyses, tested the package, and revised the manuscript. MJG contributed to study supervision. FI conceived the study and the package, contributed to the design of the benchmark analyses, wrote, and revised the manuscript, supervised the study.

## Acknowledgements

We thank Paula Weidemueller and Evangelia Petsalaki for critically reading and discussing the manuscript, and Lucia Trastulla for testing CoRe and evaluating its computational time requirements across different architectures.

## Notes

https://github.com/DepMap-Analytics/CoRe

