## Additional Files - S Figures, S table Legends, documentation for "CoRe: A robustly benchmarked R package for identifying core-fitness genes in genome-wide pooled CRISPR-Cas9 screens"

### Supplementary Figures

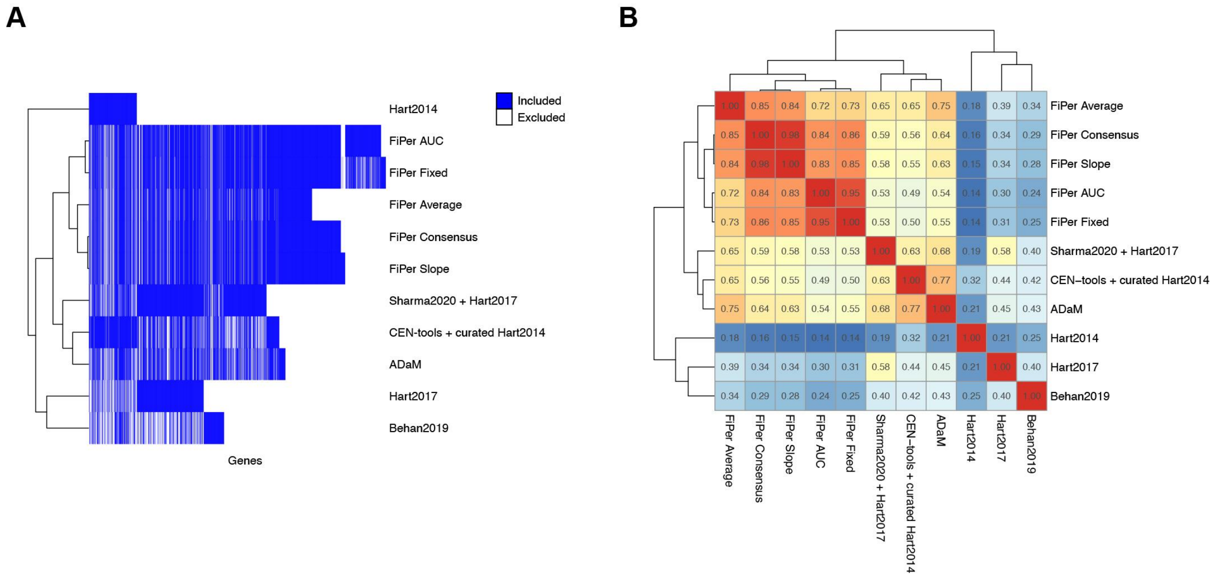


**Additional File 2 - Figure S1:** Core-fitness essential and common-essential (CFG, and CEG) sets similarity. **A.** Heatmap showing core-fitness set membership for all genes predicted as core-fitness (in the columns) by at least one method/set. **B.** Jaccard coefficient of similarity among compared core-fitness sets. The Jaccard similarity is defined as the size of the intersection divided by the size of the union of two sets.

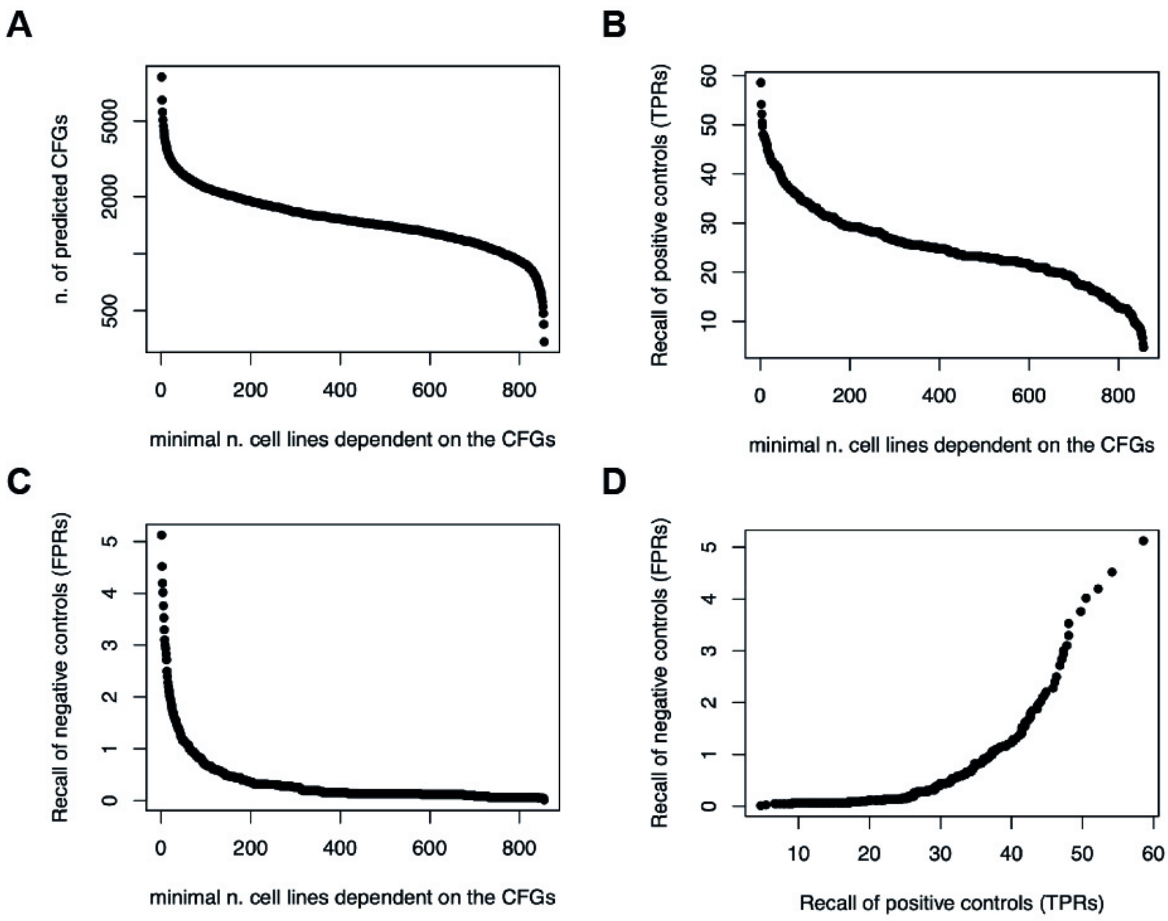


**Additional File 4 - Figure S2:** Baseline daisy model predictor (DM) performances on the DepMap dataset. **A.** Number of genes predicted as core-fitness by a baseline DM classifier (baseline core-fitness genes (CFGs)), as a function of the minimal required number of dependent cell lines, respectively y and x axis. **B.** Recall of positive controls (TPR) for each set of baseline core-fitness genes (CFGs), across all possible minimal numbers of dependent cell lines (baseline TPRs). **C.** Recall of negative controls (FPR) for each set of baseline core-fitness genes (CFGs), across all possible minimal numbers of dependent cell lines (baseline FPRs). **D.** Baseline FPR as a function of baseline TPR.

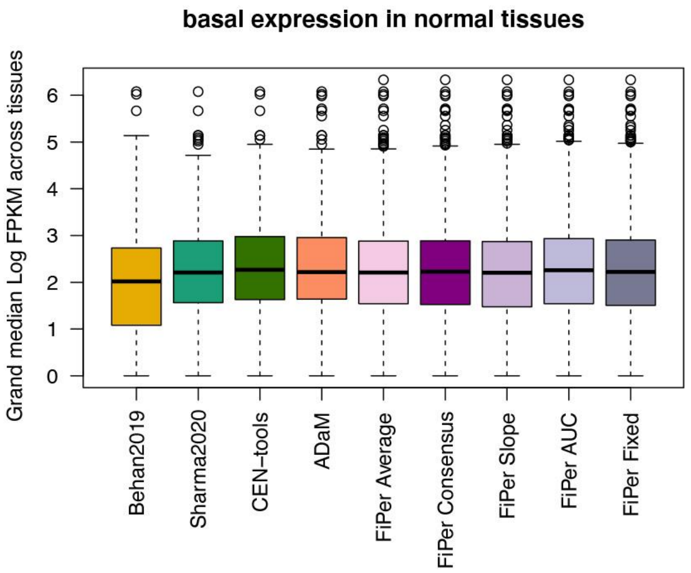


**Additional File 5 - Figure S3:** Basal expression level of predicted CFG/CEG sets in normal tissues, in terms of Fragments Per Kilobase of transcript per Million mapped reads (FPKM) extracted from the Genotype-Tissue Expression (GTEx) portal database.

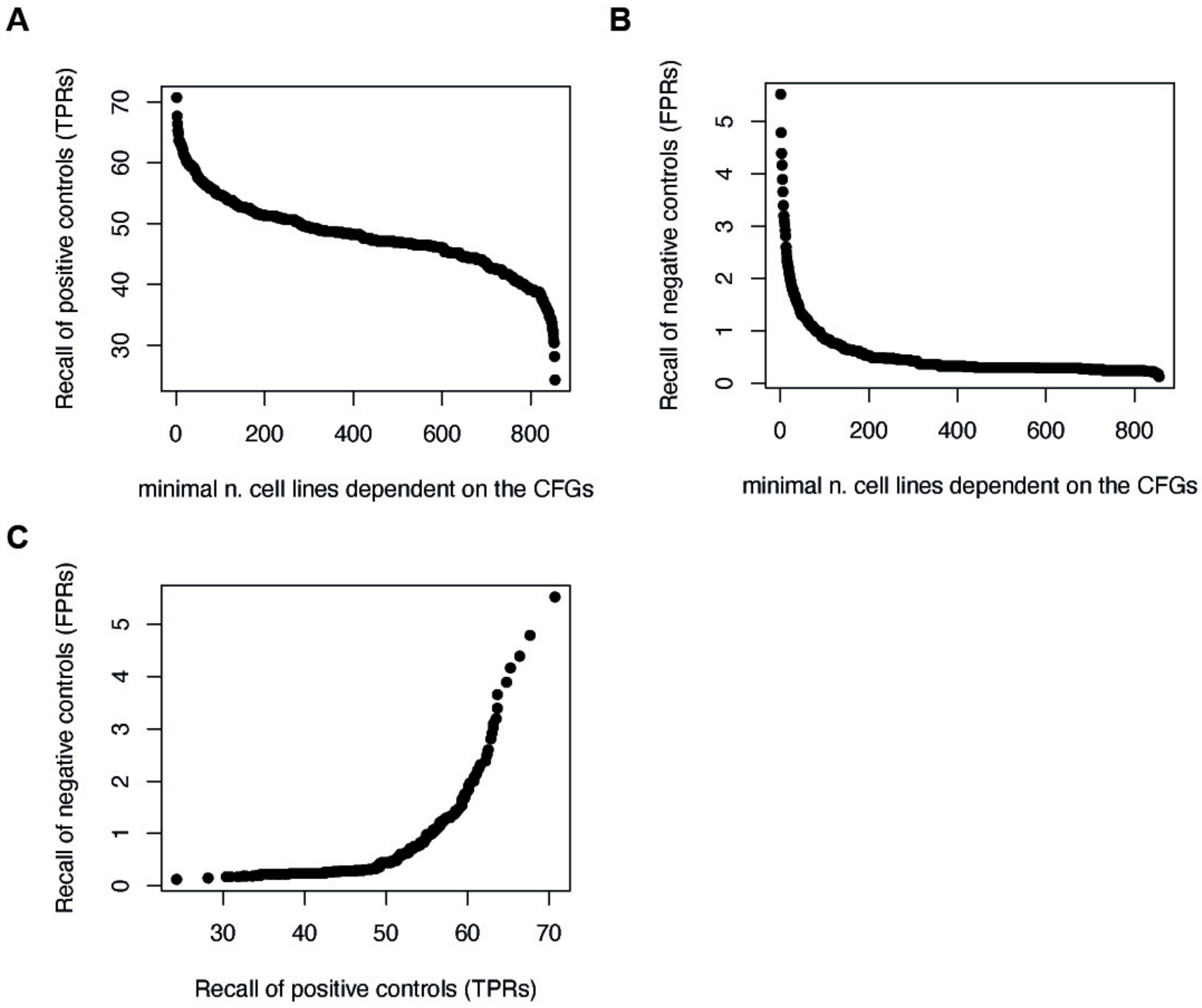

**Additional File 6 - Figure S4:** Baseline DM performances on the DepMap dataset when including genes in the training sets. **A.** Recall of positive controls (TPR) for each set of baseline core-fitness genes (CFGs) predicted by the DM as a function of the minimal required number of dependent cell lines, respectively y and x axis, across all possible minimal numbers of dependent cell lines (baseline TPRs). **B.** Recall of negative controls (FPR) for each set of baseline core-fitness genes (CFGs), across all possible minimal numbers of dependent cell lines values (baseline FPRs). **C.** Baseline FPR as a function of baseline TPR.

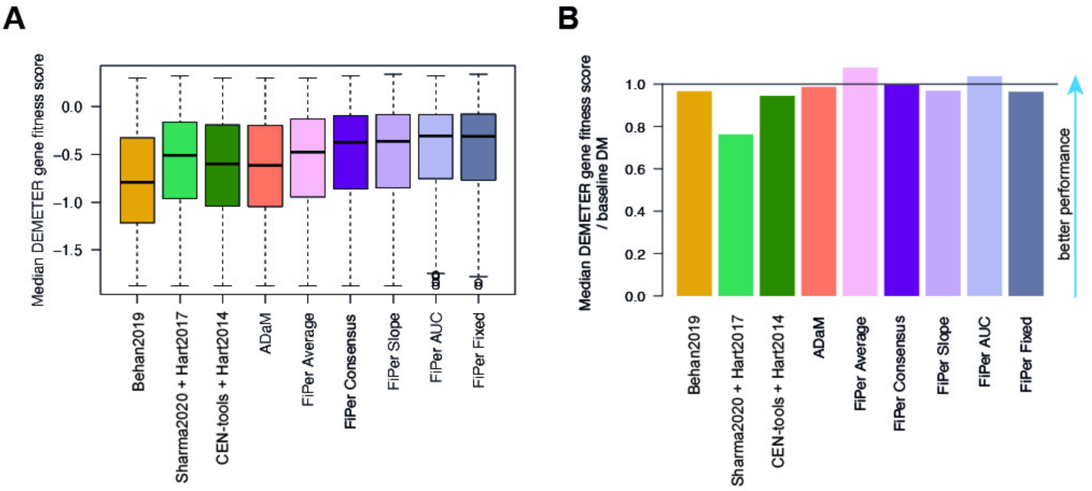


**Additional File 7 - Figure S5:** Performances' comparison considering an independent cancer dependency dataset. **A.** Fitness effect exerted by the predicted core-fitness/common-essential gene (CFG/CEG) sets using an independent RNAi based cancer dependency dataset. **B.** Ratio between the median fitness effect of each CFG set divided by the median fitness effect exerted by the baseline daisy model predictor at the observed TPRs.

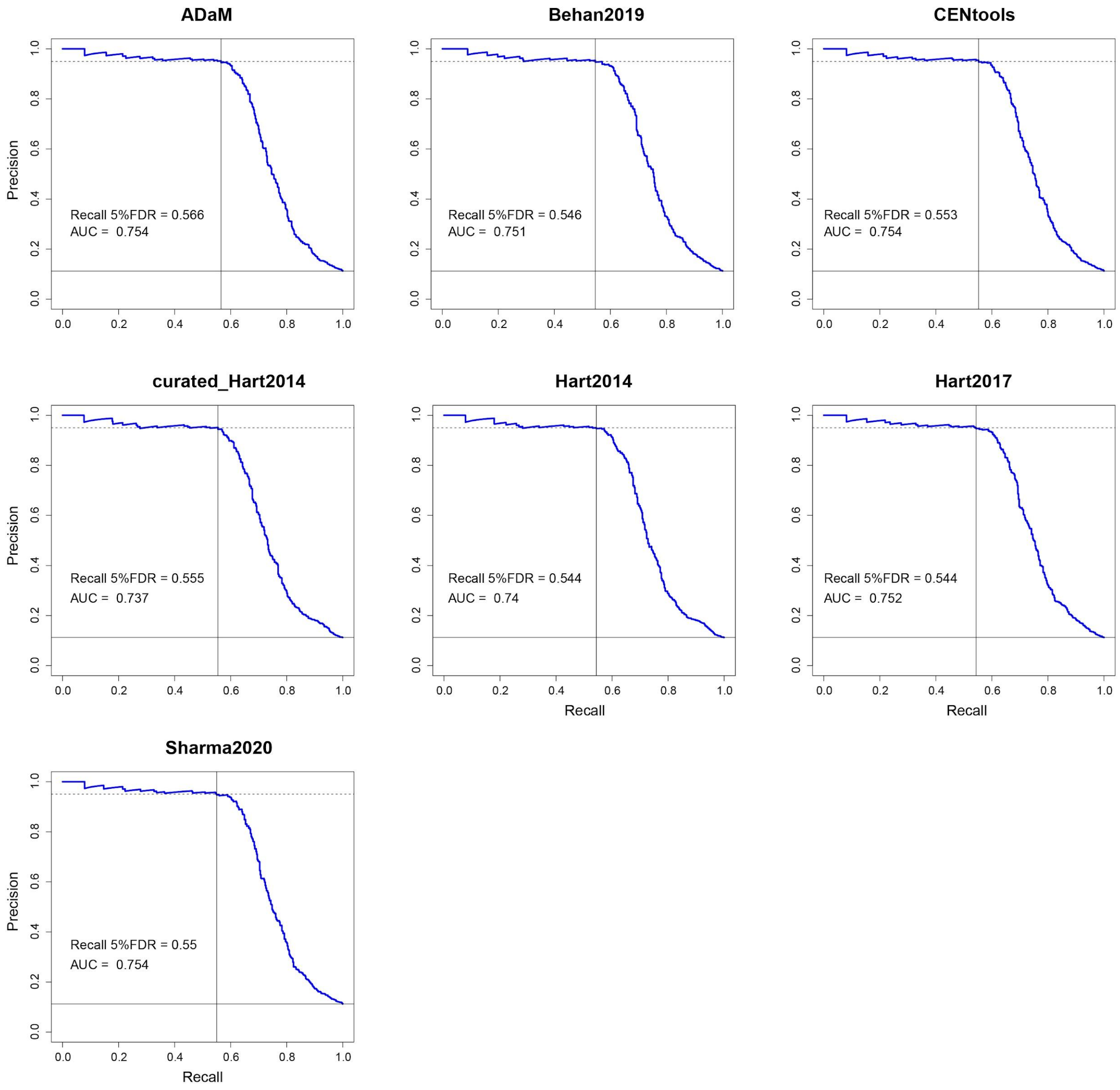


**Additional File 10 - Figure S6:** Precision-recall curves of oncogene addictions versus not-expressed oncogenes yielded by rank-based classifiers based on bayesian factors computed with BAGEL when using the compared sets of CFGs as positive training sets.

#### Legends of Supplementary Tables

**Additional File 1 - Table S1:** All compared sets of core-fitness and common-essential genes with annotations.

**Additional File 3 - Table S2:** Positive and negative control genes and their membership to training sets and DepMap datasets.

**Additional File 8 - Table S3:** Gene family enrichment analysis results.

**Additional File 9 - Table S4:** Cell line specific oncogenetic addictions, i.e. point mutated or copy number amplified oncogenes (1) and not expressed oncogenes (-1).

### Additional File 11 - Additional documentation: Running CENtools logistic regression and clustering

First, we downloaded the CENtools folder from https://gitlab.ebi.ac.uk/petsalakilab/centools/-/tree/master/CENtools. The files in the prediction subfolder were moved to the parent directory and the subfolder subsequently removed. We also removed the cluster_annotation.py. Indeed, this script should be run after the R script clustering.R, however, we don't need it for cluster object serialisation, since we saved the CENtools core essential genes as R data. Then we removed the analysis folder, since it was not used in the logistic regression.

We also downloaded the data folder from the main CENtools repository (https://gitlab.ebi.ac.uk/petsalakilab/centools/-/tree/master/) and moved it in the CENtools folder. For the purpose of our analysis, we removed unnecessary data to have a more lightweight folder:

- All data objects in the objects subfolder. Here, we have a pickle dictionary defining for each screened gene in the dataset whether it belongs to the positive training set (i.e. curated BAGEL essential) or the negative training set (i.e. curated BAGEL non-essential). To save memory, the notebook creates this object on the fly and then removes it when done.
- In the curated_data subfolder, we removed all the heavy fitness score datasets to save memory. The notebook saves the CERES_scaled_depFC.csv dataset on the fly, processed with the pipeline illustrated in the CoRe paper, and removes it when done.

We had to make some adjustments in the logistic regression script (i.e. LR.py) to make it runnable from the command line by adding a few lines of code at the end. We also commented out the figure saving instances. In this way, the script just creates and saves the pickle dictionary on the fly, executes the logistic regression model with the chosen parameters and returns only the output file of interest in a subfolder defined 'prediction_output/INTEGRATED/'. Furthermore we performed the following operations in the LR.py script:

- The code was seeded in order to guarantee reproducibility.

- In lines 44 and 332, we replaced:

*if "prediction_output" not in os.listdir(path): os.system("mkdir %s" %path + "prediction_output")*
*os.system("mkdir %s" % prediction_path + args["PROJECT"])*

With:
*if "prediction_output" not in os.listdir(path): os.mkdir(path + "prediction_output")*
*os.mkdir(prediction_path + args["PROJECT"])*
The outcome is the same (i.e. the creation of a subfolder in the designated path), however, this adjustment was needed in order for the interpreter to not throw an error.

- In line 392, we replaced:
  *bin_df = pandas.DataFrame.from_dict(bin_vector_dicts).T*

With:
*geneKeys = bin_vector_dicts.keys()*
*bin_df = numpy.array([bin_vector_dicts[i][bin_number][bin_number][j] for i in geneKeys for j in range(0, bin_number)])*
*bin_df = numpy.reshape(bin_df, (len(geneKeys), 20))*
*bin_df = pandas.DataFrame(bin_df, index=geneKeys)*
In order to save the output as a matrix that can be easily read by the R script clustering.R.

- In addition, in line 393 the typo *"bin_number"* was replaced with *"BIN_NUMBER"*.

In the paths.py script, line 4:
*path = os.getcwd() + "/venv/"*

Was replaced with:
*path = os.getcwd() + "/CENtools/"*
This adjustment makes the relative pathways available for the jupyter notebook.

Other modifications were done on the data_preparation subfolder. First, objects_.py was renamed as objects.py to make it executable from other scripts. Furthermore, we performed the following operations in the curation.py script:

- In lines 22 and 23:

*BEG = [line.strip() for line in open(curated_data_path + "Bagel_Essential_Genes.txt").readlines()]*

*BNEG = [line.strip() for line in open(curated_data_path + "Bagel_Non_Essential_Genes.txt").readlines()]*

Were replaced with:

*BEG = [line.strip() for line in open(curated_data_path + "Curated_Bagel_Essential_Genes.txt").readlines()]*

*BNEG = [line.strip() for line in open(curated_data_path + "Curated_Bagel_Non_Essential_Genes.txt").readlines()]*

This allowed to set the curated BAGEL essential and non-essential genes as default sets used in the training phase.

- In line 75:

  *integrated_cor_fc = pandas.read_csv(curated_data_path + "integrated_cor_FC_essentiality.csv", index_col=0)*

  Was replaced with:

  *integrated_cor_fc = pandas.read_csv(curated_data_path + "CERES_scaled_depFC.csv", index_col=0)*

  This allowed to set our preprocessed CERES dataset as default when performing the logistic regression.

Finally, in the R script clustering.R we commented out some parts of the code and made the function *"ClusterEssentiality"* returning just the string character vector of predicted core essential genes. Here too the code was seeded to guarantee reproducibility.
